# Lesion Quantification Toolkit: A MATLAB software tool for estimating grey matter damage and white matter disconnections in patients with focal brain lesions

**DOI:** 10.1101/2020.07.28.225771

**Authors:** Joseph C. Griffis, Nicholas V. Metcalf, Maurizio Corbetta, Gordon L. Shulman

## Abstract

Lesion studies are an important tool for cognitive neuroscientists and neurologists. However, while brain lesion studies have traditionally aimed to localize neurological symptoms to specific anatomical loci, a growing body of evidence indicates that neurological diseases such as stroke are best conceptualized as brain network disorders. While researchers in the fields of neuroscience and neurology are therefore increasingly interested in quantifying the effects of focal brain lesions on the white matter connections that form the brain’s structural connectome, few dedicated tools exist to facilitate this endeavor. Here, we present the Lesion Quantification Toolkit, a publicly available MATLAB software package for quantifying the structural impacts of focal brain lesions. The Lesion Quantification Toolkit uses atlas-based approaches to estimate parcel-level grey matter lesion loads and multiple measures of white matter disconnection severity that include tract-level disconnection measures, voxel-wise disconnection maps, and parcel-wise disconnection matrices. The toolkit also estimates lesion-induced increases in the lengths of the shortest structural paths between parcel pairs, which provide information about changes in higher-order structural network topology. We describe in detail each of the different measures produced by the toolkit, discuss their applications and considerations relevant to their use, and perform example analyses using real behavioral data collected from sub-acute stroke patients. We show that analyses performed using the different measures produced by the toolkit produce results that are highly consistent with results that have been reported in the prior literature, and we demonstrate the consistency of results obtained from analyses conducted using the different disconnection measures produced by the toolkit. We anticipate that the Lesion Quantification Toolkit will empower researchers to address research questions that would be difficult or impossible to address using traditional lesion analyses alone, and ultimately, lead to advances in our understanding of how white matter disconnections contribute to the cognitive, behavioral, and physiological consequences of focal brain lesions.

## 1. Introduction

Studies investigating the cognitive and behavioral consequences of focal brain lesions are a cornerstone of neurology and cognitive neuroscience (Rorden and Karnath, 2004). Initially, lesion studies were limited to post-mortem examinations of patients that presented with specific neurological deficits, such as speech production deficits, with the aim of localizing these deficits to lesions affecting specific anatomical loci (Berker et al., 1986). Now, modern neuroimaging techniques such as magnetic resonance imaging (MRI) and computerized tomography (CT) enable researchers to conduct large-scale lesion-behavior studies with aims that range from localizing the structural correlates of cognitive and behavioral processes (Dronkers et al., 2004; Karnath et al., 2004) to making clinical predictions about deficit severity and recovery potential (Aguilar et al., 2018; Corbetta et al., 2015; Herbet et al., 2016; Ramsey et al., 2017; Seghier et al., 2016).

Traditionally, neuroimaging-based lesion analyses such as voxel-based lesion symptom mapping (VLSM) (Bates et al., 2003) have represented brain lesions using binary voxel-wise lesion status maps that simply encode the presence vs. absence of damage to each voxel in the brain (Bates et al., 2003). These maps encode detailed information about a lesion’s spatial topography within the voxel-space coordinate system, but they do not themselves contain any explicit information about how the lesion topography maps onto the underlying brain anatomy, and group-level statistical analysis results are typically overlaid onto an anatomical T1-weighted brain template to localize effects to specific brain loci. Importantly, while binary voxel-wise lesion status representations are sufficient to characterize a lesion’s immediate spatial topography, they cannot account for the fact that this spatial topography is embedded within the dense and highly distributed network of white matter connections that together comprise the brain’s structural connectome. Accordingly, failure to account for the effects of lesions on white matter connections may obscure or distort the relationships of interest in group-level lesion analyses (Griffis et al., 2019; Pacella et al., 2019; Rudrauf et al., 2008; Thiebaut De Schotten et al., 2014).

More generally, voxel-wise binary lesion status representations are limited because they cannot directly account for the fact that different brain lesions may affect the same anatomical structures without affecting the same voxels (Griffis et al., 2019; Sperber, 2020). Anatomical structures often span many voxels, and in the case of white matter connections or distributed brain networks, the voxels associated with the structure(s) of interest may be spatially distributed across entire lobes or hemispheres. Thus, traditional lesion analyses that use binary voxel-wise lesion status representations may fail to detect relationships that involve damage to spatially distributed anatomical structures such as white matter connections, particularly when patients with damage to the same anatomical structure(s) vary with regard to the specific voxels within those structures that are lesioned (i.e. when inter-individual voxel-level lesion overlap is low).

While binary voxel-wise lesion representations provide information about a lesion’s spatial location, this information only represents the “tip of the iceberg”: a surface-level description of the lesion that is largely blind to its impact on the underlying network of white matter connections (Catani et al., 2012). Importantly, in addition to enabling large-scale lesion studies, modern neuroimaging techniques have also enabled researchers to create detailed population-scale brain atlases that delineate the boundaries of regional grey matter parcels and map the trajectories of inter-regional white matter connections (Power et al., 2011; Rojkova et al., 2016; Yeh et al., 2018; Yeo et al., 2011). These atlases can be integrated with empirical lesion data to quantify a lesion’s expected impact on specific grey matter regions and white matter connections (Carter et al., 2012; Catani et al., 2012; Foulon et al., 2018; Fridriksson et al., 2013; Griffis et al., 2020, 2019, 2017a, 2017b; Hope et al., 2015; Kuceyeski et al., 2015, 2016; Pacella et al., 2019; Thiebaut De Schotten et al., 2014), enabling advanced, anatomically-informed lesion analyses that can account for both the focal damage and distributed disconnections caused by lesions. Atlas-based lesion quantification approaches typically involve first registering a patient’s lesion to the same brain coordinate space as the desired brain atlas, and then embedding the lesion into the atlas to estimate the amount of damage/disconnection sustained by the structures of interest (Catani et al., 2012; De Schotten and Foulon, 2018; Forkel and Catani, 2018). By contextualizing the lesion in terms of *a priori* anatomical information, these approaches allow for the explicit attribution of effects to specific anatomical structures, and they may increase power in situations where patients have damage to different voxels associated with the same structures. Further, they allow for the dimensionality of the lesion data to be substantially reduced (i.e. from *n* voxels to *n* structures), which may be necessary for certain analysis approaches that can only handle a limited number of input features (Sperber et al., 2020), and which also has the advantage of reducing the number of statistical tests performed in group-level analyses and consequentially reducing the severity of multiple comparisons corrections.

The application of atlas-based approaches for quantifying white matter disconnections has enabled researchers to obtain novel insights into the distributed impacts of focal brain lesions, and has highlighted important roles of white matter disconnections in determining the cognitive, behavioral, and physiological deficits observed in brain-damaged patients (Catani et al., 2012; De Schotten and Foulon, 2018; Fox, 2018). For example, recent studies have linked deficits in broad cognitive and behavioral domains such as language (Basilakos et al., 2014; Fridriksson et al., 2013; Griffis et al., 2017a; Kümmerer et al., 2013), visuo-spatial attention (He et al., 2007; Malherbe et al., 2018; Smith et al., 2013; Thiebaut De Schotten et al., 2014), motor function (Carter et al., 2012; Feng et al., 2015; Findlater et al., 2019), and general cognition (Kuceyeski et al., 2016, 2015) to the disruption of specific white matter pathways. Other studies have employed similar approaches to test hypotheses about the effects of white matter disconnections on the structural (Foulon et al., 2018; Kuceyeski et al., 2014) and functional (Griffis et al., 2020, 2019, 2017b) properties of disconnected brain regions – research questions that would be difficult or impossible to meaningfully approach using traditional lesion analyses. For example, in recently published work, we developed and implemented novel atlas-based measures of white matter disconnection to test hypotheses about how stroke disrupts resting-state functional connectivity (Griffis et al., 2020, 2019), a functional MRI measure of inter-regional signal coherence in the resting (i.e. task-free) state that is thought to reflect ongoing neural signaling (Biswal et al., 1995; Friston et al., 2014). Similarly, other recent studies have used atlas-based lesion quantification approaches to link white matter disconnections to neurodegenerative changes in disconnected grey matter regions (Foulon et al., 2018; Kuceyeski et al., 2014). Atlas-based lesion quantification methods are therefore an increasingly important tools for brain lesion research.

However, despite the growing interest in and importance of atlas-based lesion quantification methods, relatively few publicly available software tools exist to aid researchers in implementing them in their own research (Foulon et al., 2018; Greene et al., 2019; Kuceyeski et al., 2013). Further, existing software tools tend to be relatively restricted in terms of the types of measures that they can produce, and they often rely on atlases defined based on data from relatively small samples (i.e. *N* < 100) of healthy individuals (Foulon et al., 2018; Kuceyeski et al., 2013), which may limit the generalizability of the measures that they produce. As noted above, we have recently developed and implemented novel atlas-based approaches for quantifying the severity of white matter disconnections in patients with focal brain lesions (Griffis et al., 2020, 2019). Here, we make these methods available to the broader research community by implementing them in the Lesion Quantification Toolkit, which is a publicly available MATLAB software package designed to comprehensively quantify the effects of focal brain lesions on grey matter regions and white matter connections. Unlike existing tools (Foulon et al., 2018; Greene et al., 2019; Kuceyeski et al., 2013), the Lesion Quantification Toolkit produces a comprehensive set of atlas-derived lesion measures that include measures of grey matter damage, white matter disconnection, and alterations of higher-order brain network topology. Importantly, the measures produced by the toolkit are based on population-scale (i.e. *N* > 800) atlases of grey matter parcel boundaries and white matter connection trajectories that were constructed from high-quality resting-state functional MRI and diffusion MRI data using state-of-the-art methods (Schaefer et al., 2018; Yeh et al., 2018). In the remainder of this paper, we specify the toolkit requirements and usage, provide detailed descriptions of each of the measures produced by the toolkit, and finally conclude by presenting example results obtained from real lesion-behavior analyses conducted using the measures produced by the toolkit.

## 2. Methods

### 2.1. Patient data

All example lesion and behavioral data that are reported in the paper were obtained from a sample of 132 sub-acute stroke patients that participated in the Washington University Stroke Project (Corbetta et al., 2015). Data were collected after obtaining informed consent according to procedures established by the Washington University Institutional Review Board and in accordance with the Declaration of Helsinki. Detailed descriptions of the patient sample, data collection, and data processing can be found in the original publication describing this dataset (Corbetta et al., 2015).

### 2.2. Required software

The toolkit runs in MATLAB versions 2012 and later (TheMathWorks). The toolkit requires the DSI_Studio software package in order to function correctly (freely available for download at http://dsi-studio.labsolver.org/), as MATLAB calls to DSI_Studio’s command line interface are used to create the white matter disconnection measures. All other required software packages and files (other than user-specified inputs) are included with the toolkit. These packages consist of the following: the Brain Connectivity Toolbox (Rubinov and Sporns, 2010), the Graph Theoretical Network Analysis Toolbox (Wang et al., 2015), and the matlab_nifti package (Shen, 2020). In addition, the HCP-842 diffusion MRI streamline tractography template and curated tract segmentations (Yeh et al., 2018) are also included with the toolkit, along with regional grey matter parcellations developed by Schaefer and colleagues (Schaefer et al., 2018). Additionally, while not required, external visualization packages such as MRIcroGL (https://www.nitrc.org/projects/mricrogl) or SurfIce (https://www.nitrc.org/projects/surfice) may be used to produce high-quality visualizations of the toolkit outputs.

### 2.3. Toolkit usage

The user can interact with the toolkit either via simple MATLAB scripts or via a graphical user interface (GUI) that can be accessed via the MATLAB command window. To facilitate script-based usage, example scripts for both single-subject and multi-subject (i.e. batch) processing are included with the toolkit.

User-defined inputs such as the specific regional brain parcellation and threshold parameters are saved for each run (i.e. patient) as a MATLAB structure contained within a *.mat* file, which we refer to as the “configuration file”. The information contained within the configuration file can be used to recreate the full set of outputs for each patient, thus ensuring that all outputs are easily reproducible. At any time after a patient’s lesion has been processed, summary figures illustrating the various outputs can be produced automatically using the information contained within the configuration file for that patient. Patient summary figures can be created using either the GUI or via MATLAB scripts. Detailed information about using the toolkit, including step-by-step guides for using the GUI and visualizing the outputs in external software packages, can be found in the toolkit’s accompanying User Manual (see **Supplementary Material**).

### 2.4. Toolkit inputs

The inputs provided by the user correspond to (1) a binary lesion segmentation that is registered to the Montreal Neurological Institute (MNI) template brain co-ordinate space (**Figure 1A**), and (2) a regional grey matter parcellation that is also registered to the MNI brain template space and that has identical image dimensions to the lesion segmentation (**Figure 1B**). Registration of lesions to the brain template space must be performed prior to running the toolkit, as the toolkit does not currently perform lesion-to-template registrations. Registration to template space can be accomplished using standard neuroimaging processing software such as FSL (https://fsl.fmrib.ox.ac.uk/fsl/fslwiki) or SPM (https://www.fil.ion.ucl.ac.uk/spm/), although special care should be taken to minimize registration errors introduced by the lesion (Brett et al., 2001; Griffis et al., 2017b; Ripollés et al., 2012; Seghier et al., 2008; Siegel et al., 2017).

**Figure 1.**
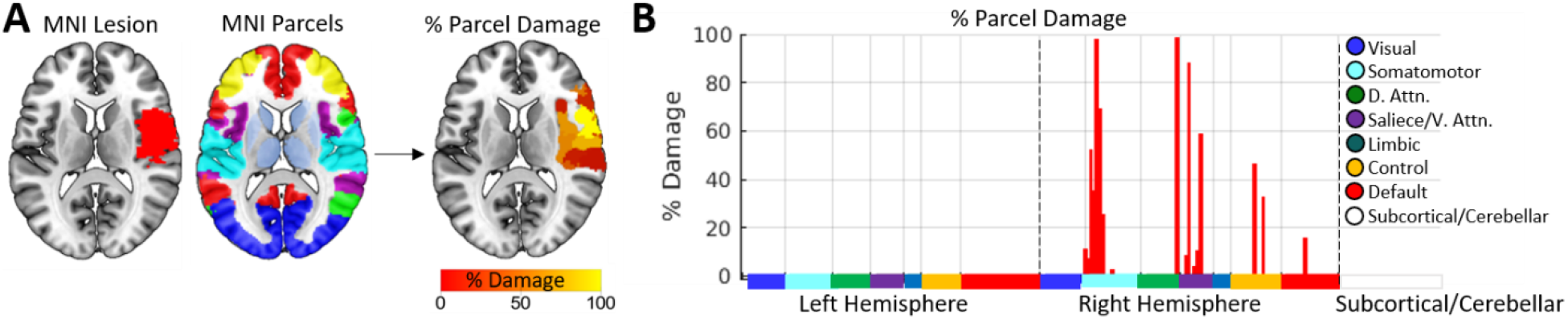
Grey matter parcel lesion loads. **A.** A lesion and brain parcellation are used to compute the percentage of voxels within each grey matter parcel that were damaged by the lesion. **B.** The percent damage sustained by each parcel is plotted as a bar graph. Parcels are split by hemisphere, and are color coded according to network assignments in the Schaefer et al., (2018) 7-network partition. Damaged parcels were located in right somatomotor, dorsal attention, salience/ventral attention, control, and default networks

While the user may select any regional brain parcellation that meets the criteria defined above, a comprehensive set of regional brain parcellations is included with the toolkit. Specifically, the toolkit includes the full set of multi-resolution cortical parcellations developed by Schaefer and colleagues (Schaefer et al., 2018), which were constructed by applying a gradient-weighted Markov random field technique to high-quality resting-state functional connectivity MRI data collected from 1489 healthy individuals. These parcellations span multiple resolutions from 100 parcels to 1000 parcels and include resting-state network assignments for each parcel. In addition, because these parcellations only include cortical areas by default, augmented versions of these parcellations that include additional subcortical and cerebellar parcels from the Automated Anatomical Labeling (AAL) atlas (Tzourio-Mazoyer et al., 2002) and a brainstem parcel from the Harvard-Oxford Subcortical atlas (https://fsl.fmrib.ox.ac.uk/fsl/fslwiki/Atlases) are also included with the toolkit to allow for complete modeling of the structural connectome, analogous to our previous work (Griffis et al., 2020, 2019). To use the included parcellations, the input lesion segmentations must have 1mm^3^ voxel dimensions and the standard MNI template image dimensions of 182×218×182. For reference, an MNI template brain volume with these dimensions is included with the toolkit, and this can be used as a registration target for external software if necessary. Alternately, the user may supply their own MNI-space brain parcellation. In any case, the lesion segmentation and brain parcellation must have identical image dimensions.

### 2.5. Toolkit outputs

#### 2.5.1. Grey matter parcel lesion loads

##### 2.5.1.1. Background

It is increasingly common for researchers to employ brain parcellations that divide the grey matter into a set of functionally or anatomically defined grey matter parcels (i.e. regions). In the context of lesion-symptom mapping, this allows researchers to utilize prior knowledge about regional boundaries to reduce the dimensionality of lesion data by summarizing it in terms of the amount of damage sustained by each grey matter parcel (Fridriksson et al., 2018; Griffis et al., 2019; Shahid et al., 2017; Sperber, 2020), often referred to as “lesion loads”. Accordingly, the toolkit includes functionality to output grey matter lesion loads for each parcel in the user-selected brain parcellation.

##### 2.5.1.2. Method

The toolkit estimates lesion loads for each grey matter parcel in the user-selected brain parcellation according to the same methods used to create the “region-based damage” measure in our previous study (Griffis et al., 2019). For a given lesion and grey matter parcellation, the toolkit computes the percentage of voxels within each grey matter parcel that overlap with the lesion, resulting in grey matter lesion loads for each grey matter parcel in the brain (**Figure 1**). The resulting parcel-wise lesion load estimates are output as both (1) a 3-D *nifti* volume where the values assigned to each grey matter parcel correspond to the percentage of voxels within that parcel that were damaged by the lesion (**Figure 1A**), and (2) a *.mat* file containing the same information stored in a 1 x *n* parcels MATLAB array (visualized in **Figure 1B**). The former can be used to visualize the severity of damage to each parcel, and the latter is formatted so that it can be easily vertically concatenated across patients to form input matrices for group-level statistical analyses.

##### 2.5.1.3. Applications and considerations

Reducing the dimensionality of lesion data by summarizing voxel-level grey matter damage in terms of parcel lesion loads contextualizes voxel-level lesion status information by incorporating *a priori* information about regional boundaries. Importantly, parcel lesion loads can be combined with measures of white matter disconnection (discussed in subsequent sections) to obtain a “complete” but relatively low-dimensional representation of a lesion’s impact on brain structure (Fridriksson et al., 2013; Griffis et al., 2019, 2017a; Pustina et al., 2017a, 2017b; Smith et al., 2013). Additionally, because the default grey matter parcellations used by the toolkit include resting-state network assignments for each grey matter parcel, parcel-level damage estimates can also easily be contextualized in terms of resting-state networks, as shown in **Figure 1B** where parcel network assignments are plotted as colored bars across the x-axis of the bar graph. The plot in **Figure 1B** shows that the lesion shown in **Figure 1A** primarily damaged parcels associated with the right somato-motor network along with parcels associated with various networks involved in attention and cognitive control.

It should be noted that parcel definitions vary across different parcellation schemes, and parcellations defined based on anatomical features (e.g. AAL) may differ substantially from those based on functional properties (e.g. Schaefer). In general, we encourage the use of functional parcellations due to the fact that they are explicitly defined to maximize regional homogeneity in terms of functional signals and/or connectivity patterns. Nonetheless, even for functional parcellations such as those included with the toolkit, parcel definitions vary (albeit typically in minor ways) across parcellation resolutions (Schaefer et al., 2018), and the appropriate parcel resolution for a given analysis will likely depend on the expected granularity of the effects of interest.

#### 2.5.2. White matter tract disconnection severities

##### 2.5.2.1. Background

The white matter connections of the human brain are often described in terms of canonical macroscale white matter tracts (Catani et al., 2002; Yeh et al., 2018) that include association pathways such as the arcuate fasciculus (AF) and superior longitudinal fasciculus (SLF), projection pathways such as the cortico-spinal tract (CST) and cortico-thalamic (CT) projections, and commissural pathways such as the corpus callosum (CC) and anterior commissure (AC). It is therefore common for researchers to measure white matter disconnections at the level of these macroscale white matter tracts (Forkel and Catani, 2018; Fridriksson et al., 2013; Griffis et al., 2019, 2017a; Hope et al., 2015; Pacella et al., 2019; Thiebaut De Schotten et al., 2014). Accordingly, the toolkit includes functionality to output white matter tract disconnection severities.

##### 2.5.2.2. Method

The toolkit estimates tract-level disconnection severities for each of 70 canonical macroscale white matter tracts (**Figure 2A**) included in the HCP-842 population-averaged streamline tractography atlas (Yeh et al., 2018), as in our previous work (Griffis et al., 2019). Detailed descriptions of the atlas construction methods can be found in the original publication by Yeh et al., (2018). Briefly, the HCP-842 atlas used by the toolkit was built using high spatial and high angular resolution diffusion MRI data collected from *N*=842 healthy Human Connectome Project participants. These data were reconstructed in the MNI template space using q-space diffeomorphic reconstruction (Yeh and Tseng, 2011), and the resulting spin distribution functions (SDFs) were averaged across all 842 individuals to estimate the normal population-level diffusion patterns. Whole-brain deterministic tractography was then performed on the population-averaged dataset using multiple turning angle thresholds to obtain 500,000 population-level streamline trajectories (i.e. estimated white matter fiber trajectories based on directional diffusion information), and the streamline trajectories were finally manually vetted and assigned to known white matter fiber tracts by a team of neuroanatomists (Yeh et al., 2018). Accordingly, the HCP-842 atlas is well-suited for use as a reference for the normal population-level white matter anatomy. We note that while the original HCP-842 streamline tractography atlas treats the corpus callosum as single tract, we have further divided the corpus callosum into five segments based on the Freesurfer corpus callosum segmentation in order to improve the interpretability of callosal disconnections, as in our previous work (Griffis et al., 2019).

**Figure 2.**
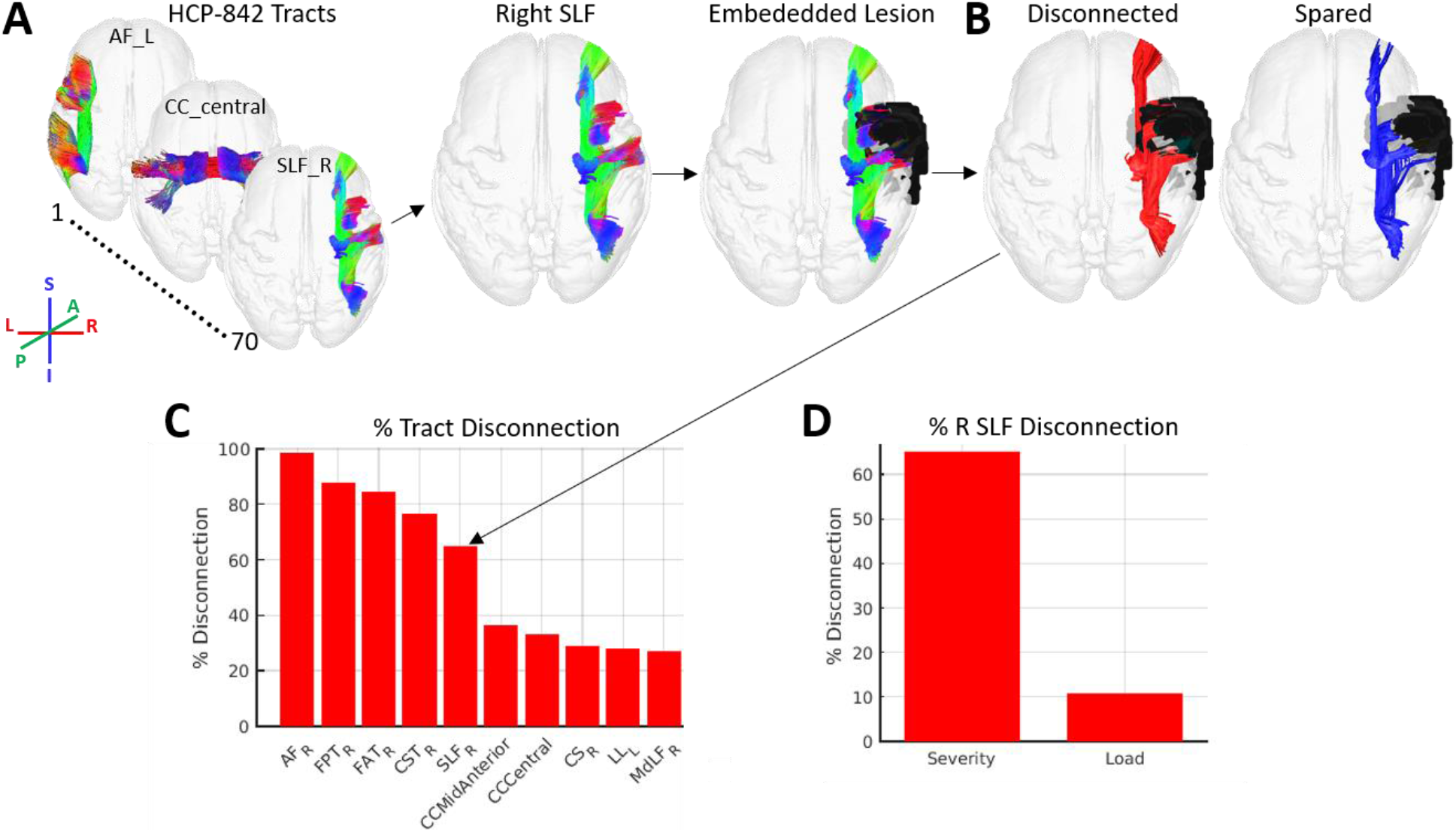
White matter tract disconnection severity. **A.** The process for estimating white matter tract disconnection severity is illustrated using the same lesion shown in **Figure 1**. For each of the 70 tracts in the HCP-842 tractography atlas (Yeh et al., 2018), the tract is loaded (i.e. right SLF) and the streamline trajectories are then intersected with the atlas-embedded lesion to identify the subset of streamlines whose trajectories intersect the volume occupied the lesion. Note: streamline trajectories are colored according to direction, as indicated by the legend. **B.** Streamlines that intersect the lesion-occupied volume are considered disconnected (red), while streamlines that do not intersect the lesion-occupied volume are considered spared (blue). **C.** The proportion of streamline trajectories that are disconnected (relative to the total number of streamline trajectories) is converted to a percentage to allow for interpretation in terms of disconnection severity. The bar graph shows the percent disconnection severities for the 10 most severely affected tracts, ordered by severity. The arrow shows the percent disconnection severity for the right SLF, which indicates that nearly 70% of the streamlines associated with the right SLF were disconnected by the lesion (see arrow). **D.** Estimates of right SLF disconnection are shown when estimated using the disconnection severity approach employed by the toolkit (x-axis: Severity) and when estimated using the lesion load approach commonly employed in the literature (x-axis: Load). Tract disconnection severity for the right SLF estimates differ substantially from tract lesion load estimates, which vastly underestimate the effect of the lesion on the right SLF.

The toolkit estimates tract disconnection severities according to the same procedures used to create the “tract-based disconnection” measure used in our previous study (Griffis et al., 2019). To estimate disconnection severity for each tract, the lesion is first embedded into the HCP-842 streamline tractography atlas as a region-of-interest (ROI) (**Figure 2A**). Then, *.trk* files containing the streamline trajectories for each of the 70 canonical white matter fiber tracts are iteratively loaded and filtered to retain only the subset of streamlines from each tract whose trajectories intersect the volume occupied by the lesion (i.e. disconnected streamlines; **Figure 2B**). For each tract, the number of disconnected streamlines is then converted to a percentage of the total number of streamlines assigned to that tract in the HCP-842 atlas, resulting in an estimate of percent disconnection severity for each tract (**Figure 2C**). The results are output as a *.mat* file containing a 1 × 70 MATLAB array containing the percentage of streamlines within each tract that are expected to be disconnected by the lesion, and a 1 × 70 MATLAB cell array containing labels for each of the 70 tracts.

##### 2.5.2.3. Applications and considerations

Like parcel lesion loads, tract disconnection measures significantly reduce the dimensionality of lesion data by describing them in terms of the disconnections incurred by a relatively small number (i.e. typically < 100) of white matter structures (Rojkova et al., 2016; Yeh et al., 2018). Importantly, as noted in the previous section, tract disconnection measures can be combined with parcel lesion load measures to obtain a complete-but-compact anatomically-informed description of the grey matter damage and white matter disconnections caused by a given lesion (Fridriksson et al., 2013; Griffis et al., 2017a; Pacella et al., 2019; Pustina et al., 2017b; Smith et al., 2013). This allows the lesion data, typically expressed in terms of several hundred thousand voxels, to be represented in terms of several hundred (or fewer) anatomical structures.

We note that the tract disconnection severity approach employed by the toolkit differs conceptually from many common approaches to estimating tract-level disconnections. Specifically, it produces an explicit measure of tract disconnection severity rather than a measure of tract lesion load or disconnection “probability” (Foulon et al., 2018; Thiebaut De Schotten et al., 2014). Analogous to parcel lesion load measures, tract lesion load measures are based on the percentage of voxels contained within each tract that overlap with the voxels contained within lesion. However, lesion load measures assume that the fundamental units composing the structure(s) of interest can be adequately summarized in terms of individual voxels. While this may be a reasonable assumption to make about the fundamental units that compose grey matter regions (i.e. populations of neuronal cell bodies), it is not a reasonable assumption to make about the fundamental units that compose white matter tracts (i.e. mathematically: streamlines; biologically: axons), since these units have directional spatial trajectories that often span multiple voxels (Hope et al., 2015). Accordingly, the fundamental elements of white matter tracts can be equivalently interrupted by a lesion anywhere along their trajectory, and damage to the same fundamental element across multiple voxels is essentially redundant in terms of its anticipated effects on signal transmission along that element. In other words, a lesion that destroys the entire length of an axon (i.e. has high lesion load on that axon) should, in principle, have essentially the same effect on that axon’s ability to carry a signal as a lesion that only bisects the axon at a single point in its trajectory (i.e. has low lesion load on that axon). Measures of tract lesion load, however, assume that the effects of the former should be more severe than the effects of the latter, and they therefore conflate the amount of voxel-wise overlap with the severity of disconnection. In principle, measures of tract disconnection severity, which account for the fact that the fundamental elements of white matter tracts have multi-voxel directional trajectories, should therefore provide a more biologically realistic measure of disconnection than measures of tract lesion load (Hope et al., 2015). Nonetheless, it should also be noted that because fiber tracts may carry multiple distinct inter-regional connections, disconnections of different portions of a given tract may still have different effects on inter-regional signal transmission, although this potential limitation can be addressed by modeling inter-regional disconnections explicitly (see next section). The discrepancy between tract disconnection severity and tract lesion load measures is illustrated in **Figure 2D**.

In addition, it is worth noting that while measures of tract disconnection “probability” are also commonly used in the literature (Foulon et al., 2018; Pacella et al., 2019; Thiebaut De Schotten et al., 2014), they are also conceptually problematic. This is because they do not actually measure the probability that a specific tract will be disconnected given a specific lesion topography (Forkel and Catani, 2018). Instead, tract disconnection “probability” measures represent the maximum (i.e. across voxels) prior probability that a streamline associated with a given tract will occupy at least one voxel that overlaps with the lesion, where the prior probabilities at each voxel are defined based on the group-level streamline visitation frequencies in a healthy reference group. Thus, a patient whose lesion overlaps with only a single voxel that has a 50% group-level tract prior probability in the healthy reference group can be assigned a 50% disconnection “probability” for the entire tract (Forkel and Catani, 2018). While this approach attempts to account for individual anatomical variability, it ultimately conflates the prior probability that a streamline associated with a given tract will be present at a given voxel in healthy individuals with the probability that a lesion affecting that voxel will disconnect the entire tract. Additionally, like the tract lesion load approach, it also ignores the fact that white matter tracts have multi-voxel directional trajectories that carry relevant information about the expected effects of a given lesion on signal transmission. While the disconnection severity approach used here ignores individual anatomical variability under the assumption that the locations and trajectories of the HCP-842 atlas tract reconstructions represent stable and conserved features of the white matter anatomy at the population level, this assumption is not unreasonable given the extremely large dataset used to construct the HCP-842 atlas (N=842) and the law of large numbers. Importantly, unlike tract lesion loads or tract disconnection “probabilities”, tract disconnection severities are conceptually aligned with principles of neural signal transmission.

#### 2.5.3. White matter disconnection maps

##### 2.5.3.1. Background

In some circumstances, it may be preferable to represent white matter disconnections in terms of their 3-dimensional spatial topographies encoded in white matter disconnection maps. For example, disconnection maps allow for the efficient visualization of the entire disconnection topography associated with a given lesion. Accordingly, the toolkit includes functionality to output 3-dimensional white matter disconnection maps.

##### 2.5.3.2. Method

The toolkit outputs 3-dimensional streamline tractography maps of the disconnections associated with a given lesion. These maps are created by embedding the lesion segmentation into the full set of streamlines aggregated over all 70 tracts in the HCP-842 streamline tractography atlas, and then filtering the streamlines to retain only the subset of streamlines that intersect the lesion (**Figure 3A**). The resulting disconnected streamline map is then output as a 3-dimensional .*trk* file that can be loaded into DSI_Studio or other tractography viewers for high-quality visualization of the disconnected streamline topographies associated with a lesion (**Figure 3A**, right image). The disconnected streamline map is also output as a 3-dimensional *nifti* file containing a tract density image (TDI) volume that represents the disconnected streamlines in voxel space (Calamante et al., 2010). This file is a raw voxel-wise disconnection map, where voxel values correspond to the densities of disconnected streamlines within each voxel. This raw voxel-wise disconnection map is then converted into a voxel-wise percent disconnection severity map where the voxel values correspond to the percentage of all of the streamlines contained within each voxel (i.e. computed from the HCP-842 streamline tractography atlas) that are expected to be disconnected by the lesion (**Figure 3B**). Optionally, spatial smoothing may be applied to the final output depending on user-specified options.

**Figure 3.**
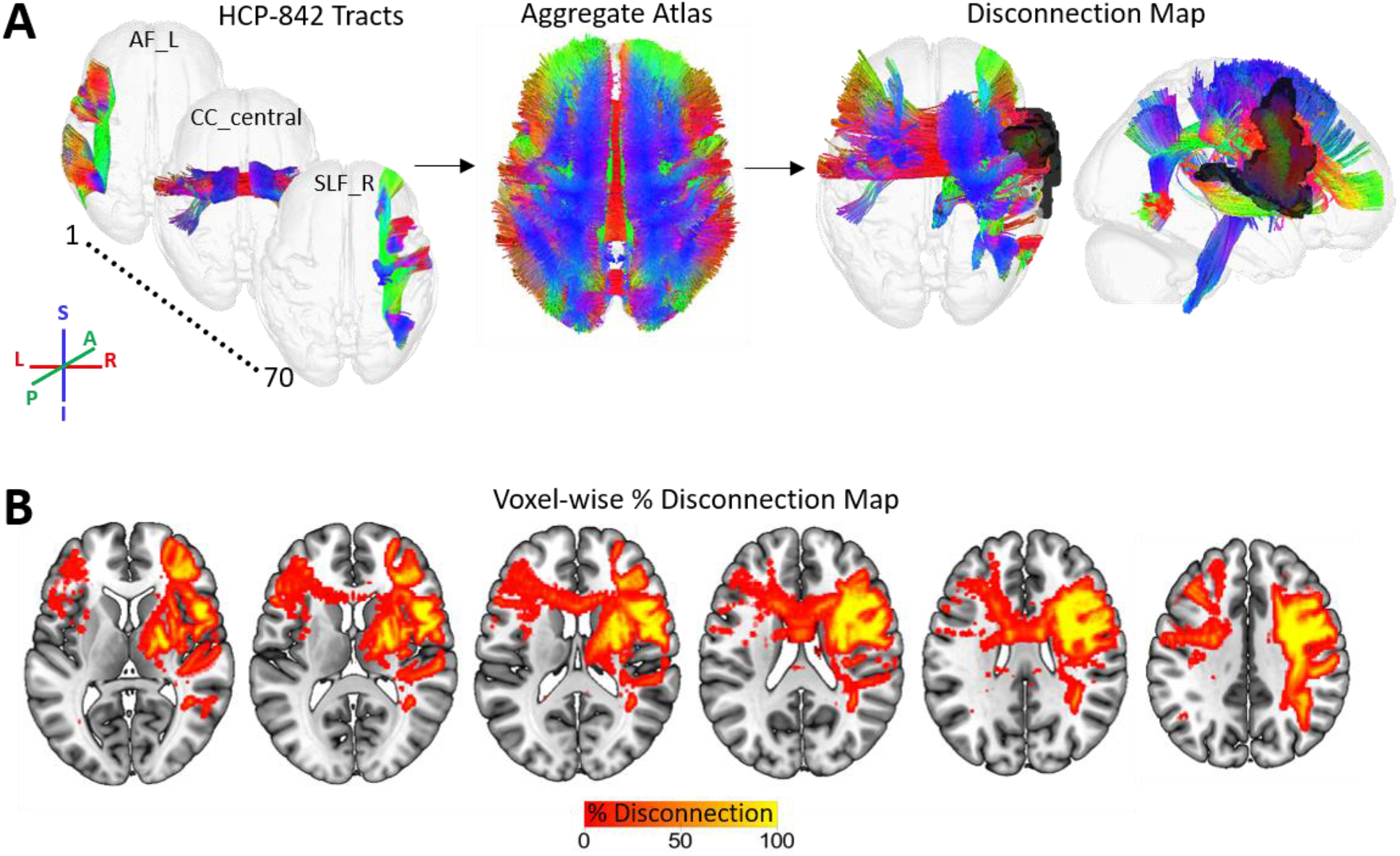
White matter disconnection maps. **A.** The process for creating white matter disconnection maps is illustrated using the same lesion shown in previous figures. Streamline trajectories for all 70 tracts from the HCP-842 streamline tractography atlas are combined into a single aggregate whole-brain streamline atlas. The lesion (black) is then embedded into the whole-brain streamline atlas and intersected with the streamline trajectories to identify the subset of streamlines whose trajectories intersect the volume occupied the lesion, producing a 3D map of disconnected streamlines. Note: streamline trajectories are colored according to direction, as indicated by the legend. **B.** The disconnected streamline map shown in (A) is output as a tract density image (TDI) volume in voxel-space and converted to a voxel-wise percent disconnection map. The map shown in (B) is smoothed with a 2mm full-width half-maximum Gaussian kernel.

##### 2.5.3.3. Applications and considerations

Map-based representations of white matter disconnections can be particularly useful for visualizing the entire pattern of white matter disconnections associated with a given lesion, including fiber endpoints in the grey matter (**Figure 3**). In addition, when white matter disconnection maps are represented in voxel space, they can be used as inputs into voxel-based analyses such as voxel-based lesion-symptom mapping or voxel-based morphometry to identify voxel-level disconnection topographies that are associated with an outcome of interest (Ashburner and Friston, 2000; Bates et al., 2003).

The percent voxel-wise disconnection severity map can be used to form inputs for traditional voxel-based analyses such as voxel-based morphometry (Calamante et al., 2010). It is important to note, however, that a given voxel may contain streamlines associated with multiple white matter pathways (Jones et al., 2013; Yeh et al., 2018), and this information is lost when the streamline maps are converted to voxel-wise TDI maps. The results of group-level analyses based on voxel-wise disconnection severity maps, which encode the expected percent reduction in streamline density for each voxel relative to the atlas due to the lesion, may therefore be best interpreted in terms of percent expected voxel-wise white matter loss. Accordingly, while voxel-wise disconnection maps can be used to infer the voxel-wise topographies of affected/implicated connections, they typically should not form the sole basis for inferences about the involvement of specific pathways due to the potentially degenerate mapping between voxel-level streamline densities and specific white matter connections.

Conceptually, the voxel-wise disconnection severity maps output by the toolkit should be more informative than the voxel-wise disconnection “probability” maps produced by other tools (Foulon et al., 2018). Similar to tract disconnection “probabilities”, disconnection “probability” maps simply quantify the probability that each voxel will contain at least one streamline that intersects the lesion based on the group-level voxel-wise streamline visitation frequencies in a healthy reference group (Foulon et al., 2018). However, since each voxel may contain streamlines associated with multiple pathways, two patients whose lesions disconnect separate white matter pathways that always intersect the same voxel in the healthy reference group could nonetheless both be assigned disconnection probabilities of 100% at that voxel. Voxel-wise disconnection probabilities are therefore difficult to meaningfully interpret.

#### 2.5.4. Parcel-wise disconnection severities

##### 2.5.4.1. Background

White matter disconnections can also be represented in terms of pair-wise disconnections between grey matter parcels. Under this representation, disconnections between parcel pairs are represented as edges between nodes (i.e. grey matter parcels) in a graph. This information can be organized as an adjacency matrix (i.e. disconnection matrix) where each cell of the matrix contains information about the severity of disconnections sustained between a pair of grey matter parcels. Disconnection matrices therefore capture the entire pattern of pair-wise white matter disconnections between grey matter parcels. Accordingly, the toolkit includes functionality to output parcel-wise disconnection matrices based on the user-selected brain parcellation.

##### 2.5.4.2. Method

Parcel-wise disconnection matrices are computed in the same way as the “region-based disconnection” measure used in our previous study (Griffis et al., 2019). Prior to estimating the parcel-wise disconnection severities for a given lesion, an atlas structural connectivity matrix is first created using the aggregate (i.e. all 70 tracts) HCP-842 streamline tractography atlas and the user-selected brain parcellation (**Figure 4A**). By default, structural connections between a parcel pair are defined as the number of atlas streamlines that bilaterally terminate within both parcels. Alternatively, the user may select a more liberal “pass-through” criterion to define parcel-wise structural connections as the number of atlas streamlines whose trajectories simply intersect a pair of parcels, although this definition may be biologically implausible and lead to interpretative difficulties. Once the atlas structural connectivity matrix has been created, the lesion is embedded into the aggregate HCP-842 streamline tractography atlas as a region-of-interest (ROI), and the atlas is filtered to retain only the subset of streamlines whose trajectories both intersect the volume occupied by the lesion (i.e. disconnected streamlines) and terminate bilaterally within a parcel pair. This results in a raw parcel-wise disconnection matrix where each entry corresponds to the number of disconnected streamlines between a parcel pair. Finally, this raw disconnection matrix is converted to a percent disconnection severity matrix relative to the atlas structural connectivity matrix (**Figure 4B**). That is, the number of disconnected streamlines between each parcel pair is converted to a percentage of the total number of streamlines connecting that parcel pair in the atlas structural connectivity matrix. The values for each cell (i.e. parcel pair) in the final percent disconnection severity matrix therefore correspond to the estimated disconnection severities for each pair of parcels. The atlas structural connectivity matrix, patient disconnected streamline matrix, and patient disconnection severity matrix are output as *n* parcels × *n* parcels MATLAB matrices contained in *.mat* files, and as .*edge* and .*node* files to allow for high-quality network visualizations using external software (**Figure 4C**).

**Figure 4.**
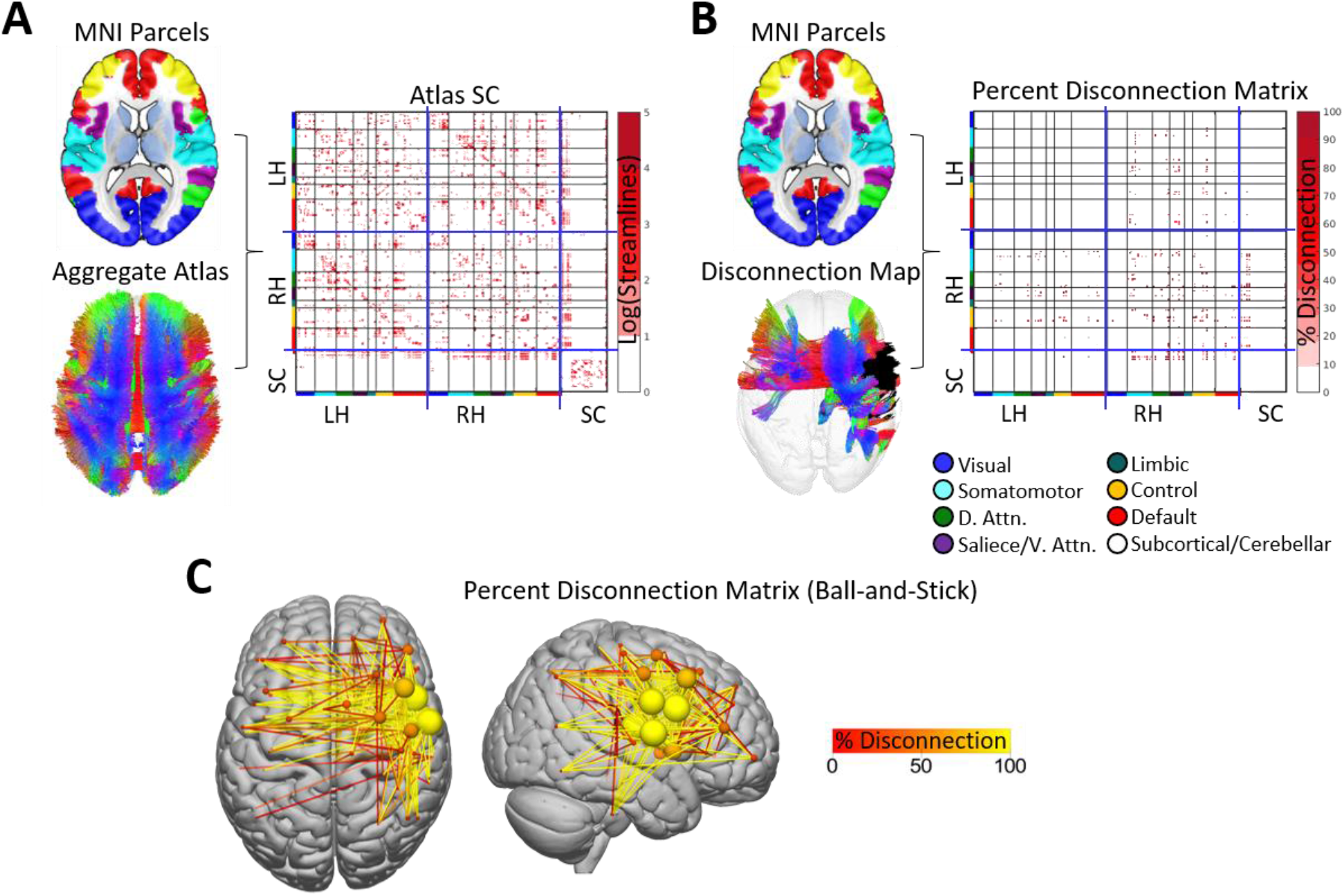
Parcel-wise disconnection matrices. **A.** The brain parcellation and aggregate HCP-842 streamline atlas (i.e. shown in **Figure 3A**) are used to create an atlas structural connectivity matrix. Each cell in the matrix encodes the number of streamlines (log-transformed for visualization) that terminate within a pair of grey matter parcels. Matrices shown are organized into blocks corresponding to left hemispheric cortical (LH), right hemispheric cortical (RH), and subcortical/cerebellar (SC) parcels (blue lines), and cortical parcels are further organized into sub-blocks corresponding to their resting-state network assignments (colored bars along matrix axes – see Figure 1; black lines). **B.** The brain parcellation, along with the disconnected streamline map (i.e. shown in **Figure 3**) and atlas structural connectome, are then used to create a percent disconnection severity matrix (i.e. disconnection matrix). **C.** The disconnection severity matrix is visualized using a ball-and-stick representation in the external SurfIce viewer. Nodes (spheres) are sized and colored according to the total disconnection severity for each node (i.e. sum of disconnection severities over all edges). Edges (lines) are colored according to disconnection severity. Connectivity matrices were computed using the Schaefer et al., (2018) 200-region parcellation.

##### 2.5.4.3. Applications and considerations

Disconnection severity matrices capture the entire pattern of pair-wise white matter disconnections between grey matter parcels, and they can be straightforwardly interpreted in terms of the expected disconnection severity between each parcel pair. Unlike other representations of white matter disconnections, disconnection matrices allow for the application of network analysis approaches based on graph theory and network science such as those implemented in the BCT toolbox included with the toolkit (Rubinov and Sporns, 2010). Accordingly, similar matrix-based representations of parcel-to-parcel white matter (dis)connectivity are commonly employed by studies investigating region-level structure-function relationships in the healthy brain (Adachi et al., 2011; Goni et al., 2014; Honey et al., 2007), and we have also recently employed parcel-wise disconnection matrices to investigate how the inter-regional functional connectivity disruptions observed after stroke relate to inter-regional white matter disconnections (Griffis et al., 2020, 2019). In addition, because they provide a parametrizable representation of the full structural (dis)connectome, matrix-based representations of white matter (dis)connectivity are also commonly employed by studies modeling the expected effects of brain lesions on the structural and functional connectomes (Alstott et al., 2009; Cabral et al., 2012; Saenger et al., 2017). Relatedly, it is worth noting that while previous lesion-connectome modeling studies have often simulated the effects of lesions on the structural connectome using parcel-wise or random connection deletion approaches, these simulation approaches cannot account for the fact that real lesions often damage the white matter and cause correlated disconnections among regions that are located within the same vascular territory (e.g. as in stroke) and/or whose connections travel in close proximity to each other (Griffis et al., 2019). To overcome these limitations, future lesion-connectome modeling studies might utilize the toolkit to more realistically simulate the disconnections expected to result from hypothetical lesions.

Like tract disconnection measures, disconnection matrices can be combined with parcel lesion load measures to obtain a “complete” representation of the grey matter damage and white matter disconnections caused by a lesion. While disconnection matrices will necessarily have higher dimensionality than tract disconnection measures, they have the advantage of allowing for a unified representation of damage and disconnections at the parcel level. Accordingly, parcel lesion load measures can also be directly integrated with disconnection matrices to differentiate between white matter disconnections involving damaged vs. undamaged grey matter parcels, and this may be important for addressing certain research questions, such as questions concerning the properties of disconnected but largely undamaged brain regions (Griffis et al., 2020).

#### 2.5.5. Parcel-wise increases in shortest structural path lengths

##### 2.5.5.1. Background

The white matter disconnections caused by focal brain lesions can disrupt communication between brain regions that are indirectly connected via a series of direct white matter connections through intermediary regions (Griffis et al., 2020; Lu et al., 2011). For example, pontine lesions disrupt a serial poly-synaptic relay connecting the somatomotor cortex with the contralateral cerebellum through an intermediate relay through the pons, and damage to the pons disrupts ongoing cortico-cerebellar functional interactions (Lu et al., 2011). Importantly, the minimum number of direct parcel-to-parcel white matter connections that must be traversed in order to establish a structural pathway between a pair of grey matter parcels, referred to as the shortest structural path length (SSPL) between those parcels, can be directly computed from a parcel-wise structural connectivity matrix (Goni et al., 2014). Intuitively, parcel pairs with direct white matter connections have SSPLs equal to 1, while parcel pairs that are only structurally connected via a series of direct connections through intermediary parcels (i.e. indirectly connected parcel pairs) have SSPLs that are equal to the number of connections in the series. Lesions that interrupt direct connections between intermediary parcels along the shortest structural path between an indirectly connected parcel pair will therefore increase the SSPL between the indirectly connected parcel pair, resulting in an “indirect disconnection” (Griffis et al., 2020). Importantly, just like directly disconnected parcel pairs (Griffis et al., 2019), indirectly disconnected parcel pairs reliably show disrupted resting-state functional connectivity (Griffis et al., 2020; Lu et al., 2011), indicating that these indirect disconnections are biologically relevant and likely disrupt inter-regional signaling. Accordingly, the toolkit includes functionality to output parcel-wise increases in SSPLs.

##### 2.5.5.2. Method

Parcel-wise SSPL increases are computed according to the same procedure used in our previous study (Griffis et al., 2020). First, the atlas structural connectivity matrix is binarized to produce a binary matrix encoding the presence vs. absence of structural connections between each parcel pair in the brain parcellation. Then, a breadth-first search algorithm (Rubinov and Sporns, 2010) is applied to the binarized atlas structural connectivity matrix to obtain an atlas SSPL matrix (**Figure 5A**). Next, a percent spared connection matrix is created for the patient by subtracting the raw (i.e. streamline count) parcel-wise disconnection matrix from the raw atlas structural connectivity matrix, and converting the resulting values to the percentage of streamlines spared (i.e. relative to the atlas structural connectivity matrix). A user-defined binarization threshold, which defines the minimum percentage of streamlines between two parcels that must be spared in order for a connection to be considered a viable link in a shortest path, is then applied to the percent spared connection matrix, producing a binary matrix where all parcel pairs have values of either 1 (i.e. spared connection) or 0 (i.e. disconnected or no direct connection). Because the patient SSPL matrix is subsequently computed based on the binarized spared connection matrix, the spared connection binarization threshold has the potential to influence analysis results. By default, the threshold value is set to 50% (i.e. at least half of the streamlines must be spared for a connection to be considered in SSPL computations), but it can be adjusted by the user, and we recommend comparing analysis results across thresholds to assess threshold dependency. After binarization, the breadth-first search algorithm is then applied to the binarized spared connection matrix to produce a patient-specific SSPL matrix where each cell encodes the SSPL between each pair of grey matter parcels in the brain parcellation. This matrix is then converted into an SSPL increase matrix by subtracting out the atlas SSPL matrix (**Figure 5B**). In the resulting SSPL increase matrix, cells corresponding to parcel pairs with preserved SSPLs have values of 0, and parcel pairs with increased SSPLs have values equal to the magnitude of the SSPL increase. If an SSPL is no longer defined between a parcel pair (i.e. no structural path between them exists, resulting an Inf value), then the SSPL between that parcel pair is assigned a value equal to 1 greater than the maximum value in the atlas SSPL matrix as in our previous work (Griffis et al., 2020). Since the SSPL increase matrix includes the magnitudes of SSPL increases for both directly and indirectly connected parcel pairs, an indirect-only SSPL increase matrix is also created by setting the SSPLs between directly connected parcel pairs equal to 0. The patient spared connection matrix, atlas SSPL matrix, patient SSPL matrix, patient SSPL increase matrix, and patient indirect-only SSPL increase matrix are output as *n* parcels × *n* parcels MATLAB matrices stored in *.mat* files, and as .*edge* and .*node* files that can be used to produce network visualizations using external software (**Figure 5C**).

**Figure 5.**
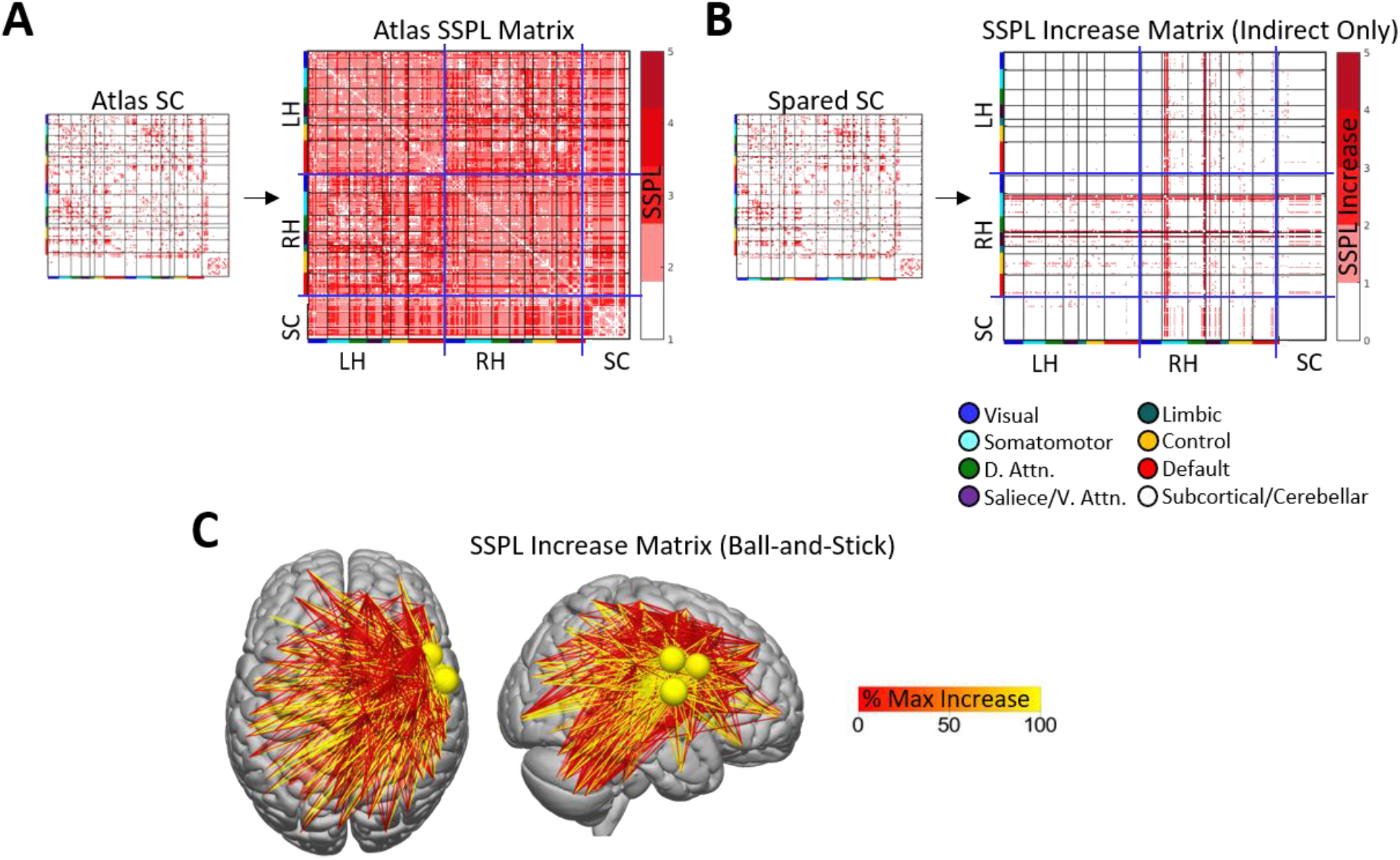
Parcel-wise SSPL increases. **A.** The atlas structural connectivity matrix is binarized and used to compute an atlas SSPL matrix. **B.** The patient spared structural connection matrix is also binarized and used along with the atlas SSPL matrix to compute an SSPL increase matrix, and parcel pairs with direct structural disconnections are removed to produce an “indirect” disconnection matrix. **C.** The “indirect” disconnections contained within the SSPL increase matrix are visualized as a ball-and-stick representation using the external SurfIce viewer. Nodes (spheres) are sized and colored according to the total SSPL increase each node. Edge (lines) are colored according to the magnitude of the SSPL increase (rescaled as % max SSPL increase across all edges). Connectivity matrices were computed using the Schaefer et al., (2018) 200-region parcellation.

##### 2.5.5.3. Applications and considerations

Parcel-wise SSPL increases provide information about how white matter disconnections alter the higher-order network topology of the structural connectome. Importantly, they provide a means for identifying “indirect” disconnections that occur due to the loss of intermediary connections along the shortest structural path between two parcels that lack direct structural connections. As noted previously, we initially developed the SSPL increase measure as an indicator of indirect disconnection in order to test hypotheses about how focal brain lesions disrupt resting-state functional connectivity after stroke (Griffis et al., 2020), and to our knowledge this measure has not otherwise been applied in the context of lesion research, although at least one other study has used a closely related measure of propagation speed to study the contribution of indirect disconnections to chronic post-stroke aphasia (Del Gaizo et al., 2017). Because we believe that this measure has the potential to enable researchers to address novel questions about how lesions disrupt the higher-order structural network topology, we felt that it was important to include this measure among the toolkit outputs.

It is worth noting that the method employed by the toolkit computes SSPLs from binarized structural connectivity matrices, as in our previous work (Griffis et al., 2020). While the breadth-first search algorithm for SSPL computation was initially chosen due to its simplicity of implementation and interpretation with respect to identifying indirectly connected parcel pairs for the purposes of our previous work (Griffis et al., 2020), other algorithms can be used to estimate SSPLs from weighted structural connectivity matrices (Goni et al., 2014). However, unlike SSPLs estimated from binary matrices, SSPLs estimated from weighted matrices cannot be straightforwardly interpreted as the number of connections along the shortest path between two regions, and we have found that SSPL increase estimates obtained using binary and weighted matrices are highly correlated (**Supplementary Figure 1**). Accordingly, the toolkit currently only includes functionality for computing SSPLs from binary matrices.

### 2.6. Damage and disconnection summary report

At any time after the damage and disconnection measures have been created for a patient, a summary report can be automatically generated from the configuration file and output files for that patient. The summary report consists of several MATLAB figure windows that each summarize different outputs from the toolkit to provide a comprehensive overview of the damage and disconnections associated with a given lesion without requiring the use of external viewing software. An example summary report is shown in **Figure 6**.

**Figure 6.**
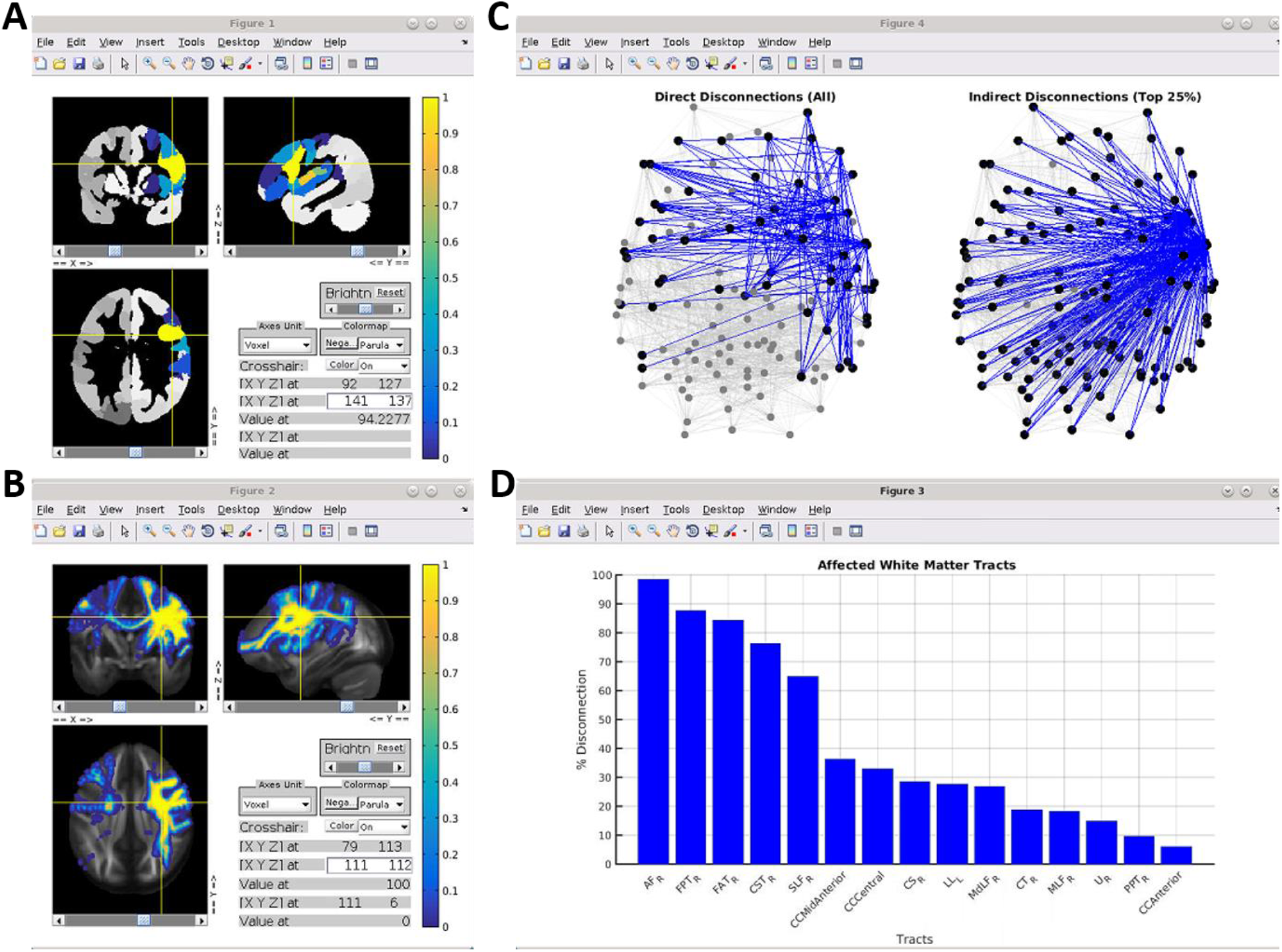
Damage and disconnection summary report. **A.** The parcel damage map is shown overlaid on three orthogonal slices from the corresponding brain parcellation in an interactive figure window. **B.** The voxel-wise disconnection severity map is shown overlaid on three orthogonal slices from the HCP-842 quantitative anisotropy volume in an interactive figure window. **C.** The direct parcel disconnection matrix (left) and indirect parcel SSPL increase matrix (right) are summarized using ball-and-stick graph brain representations in an interactive figure window. For visualization, the “indirect” disconnection matrix is thresholded to retain the top 25% of edges, although this threshold can be adjusted by the user. **D.** The tract-level disconnections are shown as a bar graph, sorted by disconnection severity. Parcel lesion loads and connectivity matrices were computed using the Schaefer et al., (2018) 200-region parcellation. Summary figures were generated using data from the same patient shown in previous figures.

### 2.7. Example analyses with real behavioral data

#### 2.7.1. Behavioral data

Finally, to illustrate the results obtained from analyses conducted using the measures output by the toolkit, we performed several example analyses using real behavioral and lesion data from the sample of sub-acute stroke patients originally described in Corbetta et al., (2015). Patients completed a battery of 42 neuropsychological tests assessing function within motor, language, attention, verbal memory, spatial memory, and visual domains (Corbetta et al., 2015). Principal component analyses (PCA) with oblique rotation were used to obtain factor scores for each domain, as described in detail by the original publication (Corbetta et al., 2015). Factor scores for the first components obtained from PCAs of the language (*n*=124), attention visual field (*n*=100), and left motor (*n=*117) domains were used for the example analyses reported here. The language factor captured general language deficits involving comprehension, naming, and reading, the attention visual field factor captured a contralesional visual field bias consistent with hemi-spatial neglect, and the left motor factor captured motor deficits affecting the left side of the body (see Table 1 in Corbetta et al., 2015). These measures were chosen because post-stroke deficits in the associated behavioral domains have been reliably linked to the disconnection of specific white matter pathways, making them well-suited for assessing the face validity of results obtained from analyses using the measures produced by the toolkit. Specifically, language deficits have been consistently linked to disconnections of the left arcuate fasciculus (AF) and left inferior fronto-occipital fasciculus (IFOF) (Catani and Mesulam, 2008; Griffis et al., 2017a; Hope et al., 2015; Ivanova et al., 2016; Kümmerer et al., 2013), hemi-spatial neglect has been linked primarily to disconnections of the right superior longitudinal fasciculus (SLF) (Shinoura et al., 2009; Thiebaut De Schotten et al., 2014; Toba et al., 2018), and contralesional motor deficits have been linked to disconnections of the ipsilesional cortico-spinal tract (CST) (Feng et al., 2015; Karnath and Rennig, 2016; Lin et al., 2018).

#### 2.7.2. Lesion data

Lesions were manually segmented on structural MRI (T1, T2, T2-FLAIR) using the ANALYZE software package (Robb and Hanson, 1991), and the lesion segmentations were reviewed by two board-certified neurologists (Maurizio Corbetta and Alex Carter). The final lesion segmentations were registered to the MNI template space using FSL’s FLIRT and FNIRT tools and resampled to 1mm^3^ voxel dimensions as described in our previous publication (Griffis et al., 2019). The MNI-registered lesion segmentations, along with the augmented (i.e. to include subcortical and cerebellar regions) 200-region cortical parcellation by Schaefer and colleagues (2018), were then input into the Lesion Quantification Toolkit to create measures of grey matter damage and white matter disconnection for each patient using the endpoint-based criterion for determining parcel-wise structural (dis)connections and a 50% spared connection threshold for computing parcel-wise SSPLs. Additional analyses were also performed using a 100% spared connection threshold as a robustness check (**Supplementary Figure 2**).

#### 2.7.3. Statistical analyses

For each behavioral measure, separate mass-univariate Pearson correlation analyses were performed using the parcel-wise grey matter lesion load, tract-wise disconnection severity, voxel-wise disconnection severity, parcel-wise disconnection severity, and parcel-wise indirect SSPL increase measures described in the previous sections. Because of the ordinal scale of the parcel-wise indirect SSPL increases, the small range (i.e. 1-5), and the observation that most SSPL increases had magnitudes of 1 (**Supplementary Figure 3**), the parcel-wise indirect SSPL increase measures were binarized prior to analysis. Supplemental analyses using the un-binarized measures nonetheless revealed generally similar results (**Supplementary Figure 2**). Additional analyses were also performed using the voxel-wise lesion maps for completeness (**Supplementary Figure 4**). All analyses were restricted to variables (i.e. parcels/tracts/voxels/connections) that were sufficiently affected (i.e. at least 10% damage/disconnection) in at least 4 patients (**Supplementary Figure 5**). To adjust for lesion volume effects on deficit severity, linear regressions were performed to regress the effects of lesion volume out of the behavioral factor scores, and the residual factor scores were used in the subsequent mass-univariate correlation analyses. False discovery rate (FDR) correction was applied to the results obtained from each analysis (Benjamini and Hochberg, 1995), and results that survived an FDR threshold of 0.05 were considered statistically significant. Due to the large number of significant effects observed for the various disconnection measures, percentile-based thresholds were applied to the FDR-corrected correlation results for display and visualization purposes (specified in **Figure 7**).

**Figure 7.**
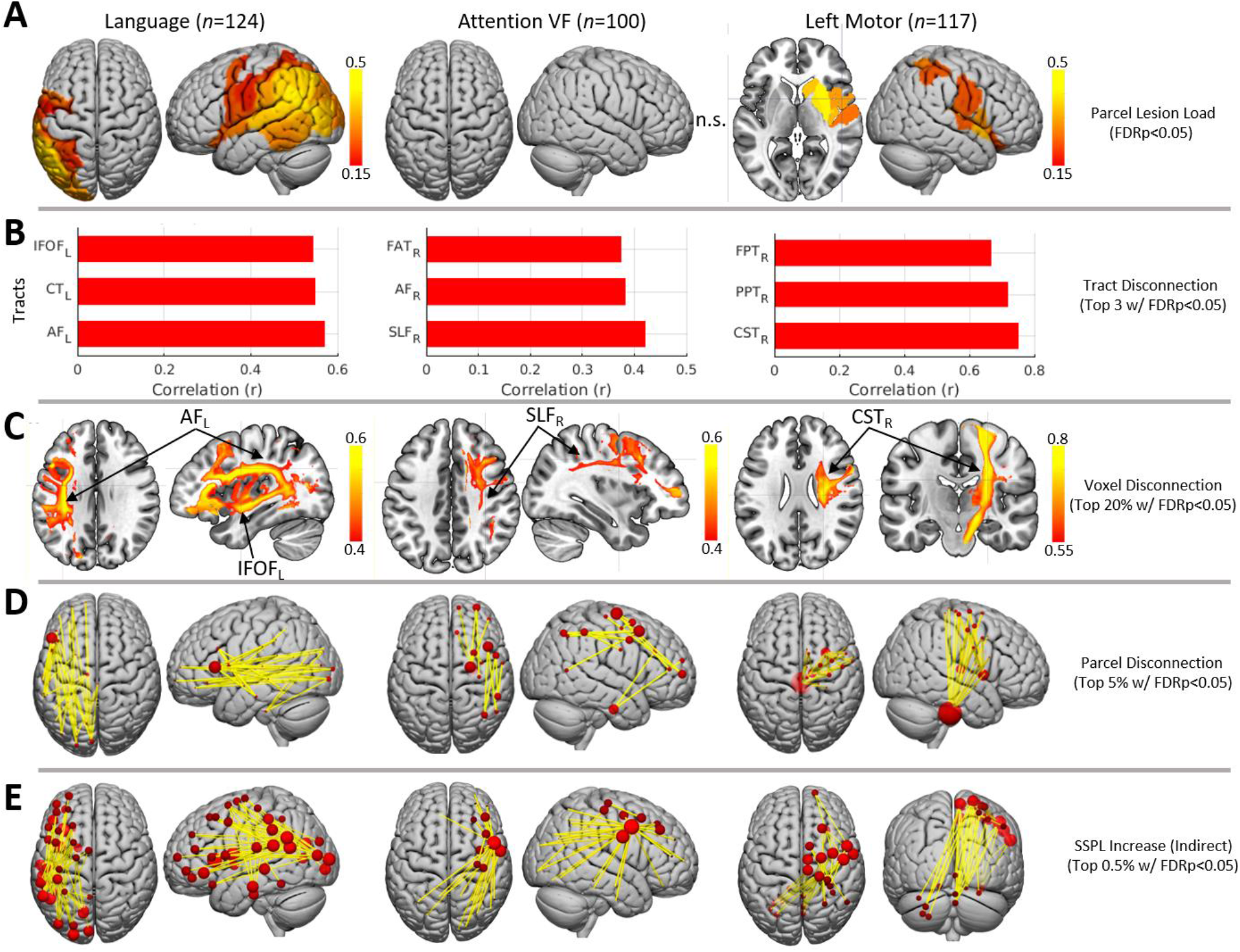
Example lesion-behavior analysis results. **A.** Significant relationships between parcel lesion loads and behavioral deficits for language (left) and left motor (right) deficits. No significant effects were identified for the attention VF measure (middle). **B.** Top 3 significant relationships between tract disconnection severities and each behavioral deficit. **C.** Top 20% of significant relationships between voxel-wise disconnection severities and each behavioral deficit. **D.** Top 5% of significant relationships between parcel-wise disconnection severities and each behavioral deficit. **E.** Top 0.5% of significant relationships between indirect parcel-wise disconnections (i.e. as indicated by SSPL increases) and each behavioral deficit. Results shown in **B-E** are restricted to strongest effects for visualization purposes. For ball-and-stick visualizations in **D-E**, ball sizes are proportional to the sum of all significant positive disconnection-deficit correlations (i.e. where disconnection correlates with more severe deficits) involving each parcel. *IFOF – inferior fronto-occipital fasciculus; CT – cortico-thalamic tract; AF – arcuate fasciculus; FAT – frontal aslant tract; SLF – superior longitudinal fasciculus; FPT – fronto-pontine tract; PPT – parieto-pointine tract; CST – cortico-spinal tract; SSPL – shortest structural path length*.

## 3. Results

### 3.1. Grey matter parcel lesion loads

Only the analyses of the language and left motor factor scores revealed significant correlations with parcel grey matter lesion loads (**Figure 7A**). The severity of language deficits most strongly correlated with lesion loads for parcels in the left posterior superior/middle temporal cortex and left temporo-parietal cortex including the supramarginal and angular gyri, while the severity of left motor deficits most strongly correlated with grey matter lesion loads in the right putamen, caudate, and insular cortex.

### 3.2. White matter tract disconnection severities

The relationships observed between tract disconnection severities and behavioral deficits were highly consistent with our expectations based on the prior literature (**Figure 7B**). The severity of language deficits was strongly correlated with the severity of disconnections sustained by the left AF and left IFOF. Language deficits were also strongly correlated with the severity of left cortico-thalamic disconnections, consistent with recent evidence implicating cortico-thalamic projections in language deficits after stroke (Griffis et al., 2017a; Mirman et al., 2015). The severity of hemi-spatial neglect, as estimated by the attention visual field factor scores, was most strongly correlated with the severity of disconnections sustained by the right SLF. Hemi-spatial neglect severity also correlated with the severity of disconnections sustained by the right AF and right frontal aslant tract (FAT), which have also been previously implicated in post-stroke visuo-spatial neglect (Carter et al., 2017; Thiebaut De Schotten et al., 2014). Left motor deficit severity was most strongly correlated with the severity of disconnections sustained by the right CST. Left motor deficit severity was also strongly correlated with the severity of disconnections sustained by the right fronto-pontine and parieto-pontine pathways, which are part of a poly-synaptic relay linking ipsilesional cortical regions to the contralesional cerebellum (Lu et al., 2011; Middleton and Strick, 2001) that has been implicated in motor deficits after stroke (den Ouden et al., 2019) and that is likely important for motor control (Stoodley et al., 2012).

### 3.3. Voxel-wise disconnection severities

The results of analyses performed using the voxel-wise disconnection severity maps were highly consistent with those obtained using the tract-wise white matter disconnection severity measures (**Figure 7C**). The thresholded maps shown in **Figure 7C** clearly show that language deficits were most strongly correlated with disconnection severities for voxels located along the trajectories of the left AF and left IFOF, while hemi-spatial neglect was most strongly correlated with disconnection severities for voxels located along the trajectory of the right SLF, and left motor deficits were most strongly correlated with disconnection severities for voxels located along the trajectory of the right CST (**Figure 7C**, third row).

### 3.4. Parcel-wise disconnection severities

Highly consistent results were also obtained from analyses performed using the parcel-wise disconnection severity matrices (**Figure 7D**). The severity of language deficits was most strongly correlated with the severity of white matter disconnections between parcels in left inferior frontal cortex and parcels in left temporal, parietal, and occipital cortices, consistent with disconnections of the left AF and left IFOF. The severity of hemi-spatial neglect was most strongly correlated with the severity of white matter disconnections between parcels in right lateral pre-frontal cortices and parcels in right temporal and parietal cortices, consistent with disconnections of the right SLF and right AF. The severity of left motor deficits was most strongly correlated with the severity of white matter disconnections between parcels in right somatomotor cortex and parcels in the right thalamus, putamen, and the brainstem, consistent with disconnections of the right CST and right cortico-thalamic/cortico-striatal projections.

### 3.5. Parcel-wise indirect disconnections identified by SSPL increases

Finally, analyses revealed significant correlations between behavioral deficits and indirect parcel-wise disconnections (i.e. as identified by SSPL increases between indirectly connected parcel pairs) for all three behavioral measures (**Figure 7E**). The severity of language deficits was most strongly correlated with indirect disconnections between left temporal/temporo-parietal/temporo-occipital parcels and parcels distributed throughout the left frontal lobe, presumably reflecting disruptions of indirect pathways connecting these parcels with distributed frontal regions via direct connections to the left inferior frontal gyrus (**Figure 7D**) through canonical fiber tracts such as the left AF and IFOF (**Figure 7B-C**). This interpretation is consistent with models that posit the left IFG as a cortical hub in frontal cortex that directly interfaces with posterior supramodal convergence zones to select and retrieve information (Binder and Desai, 2011) that may then be passed to other frontal regions such as pre-motor cortex (Basilakos et al., 2017).

The severity of hemi-spatial neglect was most strongly correlated with indirect disconnections between a parcel located near the right inferior frontal junction (IFJ) (Muhle-Karbe et al., 2015) associated with the ventral attention/salience networks (i.e. per the Schaefer et al., 2018 7-network partition) and parcels distributed throughout the right frontal, right temporal, right parietal, and right occipital lobes (**Figure 7E**, middle), suggesting that the direct disconnections associated with spatial neglect (**Figure 7B-D**, middle) tend to disrupt indirect pathways between the right IFJ and parcels distributed throughout the right hemisphere. Interestingly, this finding is consistent with previous work suggesting that this region may integrate information between the ventral and dorsal attention systems and implicating its long-range functional interactions in the pathogenesis of spatial neglect (Asplund et al., 2010; He et al., 2007). Strong correlations were also observed between indirect disconnections among right frontal regions and between right frontal regions and the left parietal lobe.

Finally, the severity of left motor deficits was most strongly correlated with indirect disconnections between parcels in right somatomotor cortex and parcels in the left/medial cerebellum (**Figure 7C**, bottom row), consistent with disruptions of the poly-synaptic cortico-pontine-cerebellar pathway by the direct disconnection of cortico-pontine projections (Lu et al., 2011). Left motor deficits were also correlated with indirect disconnections between right somato-motor cortex and parcels in right pre-frontal cortex.

## 4. Discussion

Over the past two decades, the development and implementation of advanced techniques for non-invasively measuring brain connectivity has revolutionized the fields of neuroscience and neurology by shifting their focus towards understanding how cognition, behavior, and clinical symptoms relate to distributed brain networks and their connections. Accordingly, researchers in these fields are increasingly interested in quantifying the effects of focal brain lesions on the brain’s structural connectome. To address this need, we developed the Lesion Quantification Toolkit, a publicly available MATLAB software package designed to comprehensively characterize the grey matter damage and white matter disconnections associated with pathologies such as brain tumors, traumatic brain injuries, and stroke. Here, we presented the toolkit and provided detailed descriptions of the different outputs that it creates. In addition, we demonstrated the utility of the measures produced by the toolkit by incorporating them into example analyses of real behavioral data obtained from a large sample of sub-acute stroke patients. We anticipate that this toolkit will constitute an important resource that facilitates the integration of white matter disconnection measures into studies of patients with focal brain lesions.

### 4.1. Comparisons with other software tools

As noted in the Introduction, relatively few dedicated tools exist to support researchers in quantifying the structural impacts of focal brain lesions. While all existing tools employ principally similar atlas-based approaches to quantifying lesion-induced white matter disconnections, the Lesion Quantification Toolkit offers several advantages over existing tools.

First and foremost, the Lesion Quantification Toolkit produces a comprehensive set of measures that include parcel-wise grey matter lesion loads and multiple measures of white matter disconnection that include tract-wise, voxel-wise, and parcel-wise disconnection severities. Importantly, it also estimates parcel-wise SSPL increases that can be leveraged to obtain insights into the “indirect” disconnections caused by a lesion (e.g. **Figure 6, Figure 7** – bottom row). This is in contrast to other existing tools that typically produce only one or two measures.

For example, the Network Modification (NeMo) Tool, one of the first tools developed for quantifying white matter disconnections in patients with focal brain lesions, only produces two measures for each lesion input: parcel-level scores summarizing the total (i.e. across all pair-wise connections) change in connectivity for each grey matter parcel, and whole-brain summary statistics summarizing changes in global network topology (Kuceyeski et al., 2013). Importantly, neither of these summary measures provide information about the specific white matter connections (i.e. fiber tracts, parcel-wise connections) that are affected by a lesion. Further, because the NeMo Tool does not produce any measures of parcel grey matter lesion loads, it cannot be determined whether the connectivity change scores obtained for a given parcel reflect the effects of direct damage to that parcel or the effects of damage to that parcel’s white matter connections. While the Lesion Quantification Toolkit does not directly output global network summary statistics, these measures can be easily computed from the parcel-wise disconnection, spared connection, and SSPL measures using functions implemented in the included Brain Connectivity Toolbox (Rubinov and Sporns, 2010).

Similarly, another recently developed tool, the Brain Connectivity and Behavior (BCB) Toolkit, only produces measures of tract-wise lesion loads, tract-wise disconnection “probabilities”, and voxel-wise disconnection “probabilities” (Foulon et al., 2018). However, as discussed previously in the Methods, lesion load measures are inappropriate for measuring white matter disconnections because they ignore the multi-voxel directional trajectories of white matter fibers and their streamline representations and inappropriately assume that greater overlaps between the lesion- and tract-occupied volumes will necessarily translate to more severe disconnections (**Figure 2D**). Similarly, the tract disconnection “probability” measures produced by the BCB toolkit do not actually reflect the probability that a given white matter tract is disconnected, but instead reflect the maximum prior probability that a streamline associated with a given tract in a healthy reference group will be present within at least one lesioned voxel (Forkel and Catani, 2018). In contrast, the white matter disconnection measures produced by the Lesion Quantification Toolkit are uniformly expressed in terms of disconnection severities, which are conceptually sound and can be interpreted in straightforward and biologically relevant terms. While one other recently developed tool based on shortest path tractography does measure white matter disconnections in terms of disconnection severity (which it refers to as “connectivity loss”), this tool is still relatively limited in scope, as it only produces parcel-wise disconnection matrices (Greene et al., 2019). By outputting multiple complementary measures of white matter disconnection severity, the Lesion Quantification Toolkit allows the user to select the most appropriate measure(s) to address the question of interest and/or to assess the consistency of results across multiple complementary perspectives (**Figure 7**).

Another advantage of the Lesion Quantification Toolkit is that it computes white matter disconnection measures based on the population-scale HCP-842 streamline tractography atlas. This atlas was constructed from group-averaged high spatial and high angular resolution diffusion MRI data from 842 healthy individuals, and the final streamline reconstructions were expert-vetted to improve anatomical fidelity and reduce the prevalence of false positive streamlines (Yeh et al., 2018). In contrast, the tractography atlases used by other existing methods are based on atlases constructed from much smaller diffusion MRI datasets (NeMo: *N*=70; BCB toolkit: *N*=10; Shortest path tractography: *N*=200), and it is unclear whether the underlying data used by other tools have been explicitly vetted to maximize correspondence with the known white matter anatomy, which may influence the quality of the measures (**Supplementary Figure 6**). This may be particularly important given the high prevalence of false positive connections in diffusion MRI tractography analyses (Maier-Hein et al., 2017). In addition, whereas the atlases utilized by other tools consist of multiple, separate, subject-level streamline sets that were each constructed from single-subject diffusion MRI data, the HCP-842 atlas consists of only a single group-level streamline set that was constructed from population-averaged diffusion data (i.e. averaged over all 842 subjects) in the MNI template space. While the HCP-842 atlas (and subsequently, any measure derived from it) does not explicitly account for inter-individual anatomical variability, it should nonetheless represent the most stable features of the population-level white matter anatomy due to large sample averaging of high-quality data (Yeh et al., 2018).

### 4.2. Considerations related to atlas-derived lesion measures

As with other similar tools, all of the measures produced by the Lesion Quantification Toolkit are necessarily atlas-derived measures, and so they cannot capture patient-specific anatomical features (e.g. variations in the spared anatomy), which may be relevant for explaining variability in outcomes across patients with similar lesions (Forkel et al., 2014; Forkel and Catani, 2018). Nonetheless, atlas-based measures are advantageous in the sense that they reduce the potential for confounding influences from sources such as inter-individual variation in diffusion MRI data quality and/or data reconstruction, which may be particularly relevant for studies of patients with gross anatomical abnormalities such as brain lesions (Gleichgerrcht et al., 2017; Jones et al., 2013).

Atlas-derived measures of damage and disconnection also have the desirable property of being defined relative to a common reference, which makes them directly comparable across individuals and even across independent studies. Notably, we have previously used the different measures that are produced by the toolkit with great success in our recent studies on the structural bases of functional connectivity disruptions caused by stroke (Griffis et al., 2020, 2019), and similar atlas-derived measures have been successfully employed to investigate the structural correlates of deficits in various behavioral domains including attention (Smith et al., 2013; Thiebaut De Schotten et al., 2014), language (Fridriksson et al., 2018; Griffis et al., 2017a, 2017b; Hope et al., 2015), and general cognition and daily living (Kuceyeski et al., 2016, 2015). The results of the example analyses presented here also align closely with results reported in the prior literature and provide evidence to support the face validity of the measures produced by the toolkit (**Figure 7**). Importantly, the results of these analyses demonstrate that the different white matter disconnection measures provide complementary perspectives that nonetheless yield consistent conclusions about the white matter disconnections implicated in different behavioral deficits (**Figure 7**).

It nonetheless remains unclear whether or not atlas-derived disconnection measures can consistently provide additional power to predict outcomes beyond that provided by traditional lesion measures. While some studies have reported that white matter disconnection measures are superior to traditional lesion measures for explaining behavioral and physiological disruptions in patients with focal brain lesions (Griffis et al., 2019; Kuceyeski et al., 2016, 2015; Pacella et al., 2019), other studies have found little-to-no difference in performance between lesion-based vs. disconnection-based models (Hope et al., 2018; Salvalaggio et al., 2020). Several factors likely contribute to these discrepant results across studies, including variability in the sample sizes and lesion characteristics of the samples under study, variability in how disconnection measures are defined, and variability in the degree to which the outcomes of interest actually depend directly on distributed white matter disconnections vs. focal damage to specific structures. We briefly explore these possibilities below.

For example, it may be possible for machine learning algorithms to recover implicit disconnection information when applied to voxel-level lesion data from extremely large patient samples if the group-level lesion coverage, voxel-level lesion frequencies, and inter-patient lesion variability is sufficiently high. This, along with the very coarse nature of the parcellation (i.e. AAL) employed, could potentially explain why Hope and colleagues (2018; *N=*818) did not identify substantial differences in predictive power between damage-based and disconnection-based models of chronic language deficits. It is also possible that certain types of disconnection information are less informative than others. For example, Salvalaggio and colleagues (2020) found only minor differences in the predictive power of behavioral deficit models based on voxel-wise lesion maps vs. voxel-wise disconnection “probability” maps. Speculatively, it is possible that this might reflect the loss of connection-level information incurred by representing disconnections as binary voxel-level streamline visitation maps, especially given that the maps in question conflate voxel-level streamline visitation probabilities with disconnection probabilities (see section 2.5.3) and may feature biologically implausible disconnections (**Supplementary Figure 6**). Finally, different outcomes may differ in the extent to which they actually directly depend on white matter disconnections. For example, there is strong *a priori* evidence that functional connectivity, at least in part, depends directly on the underlying structural connections through the white matter (Adachi et al., 2011; Goni et al., 2014; Johnston et al., 2008; O’Reilly et al., 2013; Roland et al., 2017; Van Den Heuvel et al., 2009), but the degree to which complex cognitive/behavioral processes depend directly on white matter structural connections is less clear. This could also potentially explain why we found white matter disconnection measures to consistently outperform parcel lesion load and voxel-wise lesion status measures for explaining variability in the severity of functional connectivity disruptions caused by stroke (Griffis et al., 2019), while other studies focusing on behavioral outcomes have not identified such consistent advantages (e.g. Hope et al., 2018; Salvalaggio et al., 2020; but see also Kuceyeski et al., 2015,2016 and Pacella et al., 2019). Nonetheless, the relative utility of traditional lesion measures vs. atlas-derived white matter disconnection measures for predicting individual clinical outcomes remains an important topic of ongoing study, and more dedicated work on this topic is necessary to ultimately determine if and/or when disconnection measures will provide additional explanatory and/or predictive power. By making this toolkit publicly available, we aim to facilitate further research in this domain.

## 5. Conclusions

The Lesion Quantification Toolkit is, to our knowledge, the most comprehensive software tool for atlas-based quantification of grey matter damage and white matter disconnections in patients with focal brain lesions. Accordingly, we anticipate that it will constitute a valuable tool that will empower researchers to obtain novel insights into how both the focal damage and distributed disconnections caused by focal brain lesions relate to cognition, behavior, and brain function.

## Supporting information

User Manual

Lesion Quantification Toolkit Software

## Data and Software Availability

The Lesion Quantification Toolkit software will be made publicly available for download upon publication and is also available via request to the corresponding author.

Lesion and behavioral data can be accessed at http://cnda.wustl.edu/app/template/Login.vm.

## Acknowledgments

Funding: R01 NS095741 and R01 HD061117 to M.C.

Data were provided [in part] by the Human Connectome Project, WU-Minn Consortium (Principal Investigators: David Van Essen and Kamil Ugurbil; 1U54MH091657) funded by the 16 NIH Institutes and Centers that support the NIH Blueprint for Neuroscience Research; and by the McDonnell Center for Systems Neuroscience at Washington University.

We thank Alessandro Salvalaggio for providing the disconnection probability maps shown in Supplementary Figure 6.

## Author Contributions

J.G designed and wrote the software and wrote the paper. J.G and N.M. performed data processing, analyses, testing. J.G., G.S., and M.C. edited the paper. G.S. and M.C. contributed data and other resources.

