## Supplementary Figures for "Lesion Quantification Toolkit: A MATLAB software tool for estimating grey matter damage and white matter disconnections in patients with focal brain lesions"

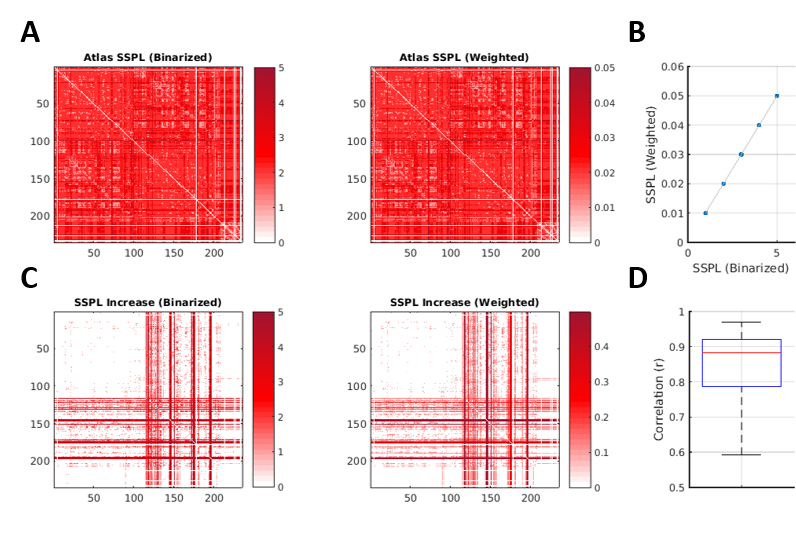

**Supplementary Figure 1.** Comparison of SSPLs computed from binarized vs. weighted structural connectivity matrices. **A.** Atlas SSPL matrices computed from binarized (left) and weighted (right) atlas structural connectivity matrices. In this context, structural connectivity weights were defined as the percentage of each connection that was spared, since this allows for intuitive and straightforward weighting of the patient structural connectivity matrices. Since patient SSPL matrices must be compared to the atlas SSPL matrix to compute SSPL increases, the atlas structural connectivity weights were defined in the same way, and all atlas connections were assigned weights equal to 100%. Accordingly, the atlas SSPL matrices computed from the binarized (left) and weighted (SC) matrices are identical aside from scaling. **B.** The scatterplot shows that the atlas SSPL matrices computed from the binarized (x-axis) and weighted (y-axis) structural connectivity matrices are linearly equivalent. **C.** Example patient SSPL increase matrices computed from binarized (left) and weighted (right) patient structural connectivity matrices. Values in the right matrix were square-root transformed to reduce the right skewness of the values and improve visualization of similarities to the left matrix. **D.** Spearman correlations were computed between SSPL increase matrices obtained from binarized and weighted spared connectivity matrices for each of 132 patients. The boxplot summarizes the distribution of correlations across patients. In general, SSPL increase matrices computed from weighted vs. binary spared connectivity matrices were highly similar across patients.

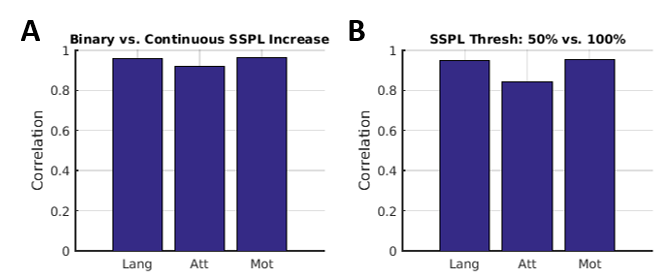

**Supplementary Figure 2.** Similarity of behavioral relationships with SSPLs using different SSPL criteria. **A.** Behavioral correlation analyses were performed using both binarized (i.e. all parcel pairs with SSPL increases were coded as 1, and all others were coded as 0) and continuous SSPL (i.e. raw SSPL increase magnitudes in steps of 1 to 5) increase matrices, and the resulting correlation matrices were correlated with each other to assess the similarity between results obtained using each method. Results obtained using each method were highly correlated for each behavioral measure, indicating that the overall pattern of results was highly similar regardless of which method was used. These analyses both used percent spared connection binarization thresholds of 50%. **B.** Behavioral correlation analyses were performed using different percent spared connection binarization thresholds to compute the patient SSPL methods, and the resulting correlation matrices were again correlated with each other. For both analyses, SSPL increase matrices were binarized so that all parcel pairs with SSPL increases had values of 1, and all others had values of 0. Results obtained using the 50% and 100% spared connection binarization thresholds were highly similar, again indicating that the overall pattern of results was highly similar regardless of which threshold was used.

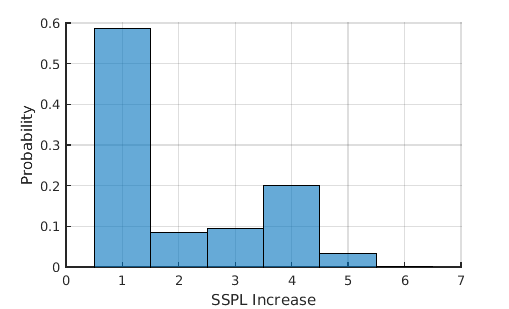

**Supplementary Figure 3.** Group-level SSPL increases. The histogram shows the distribution of non-zero SSPL increases across all 132 patients, computed using the binarized spared connection matrices and using a 50% spared connection threshold. Nearly 60% of SSPL increases had magnitudes equal to 1.

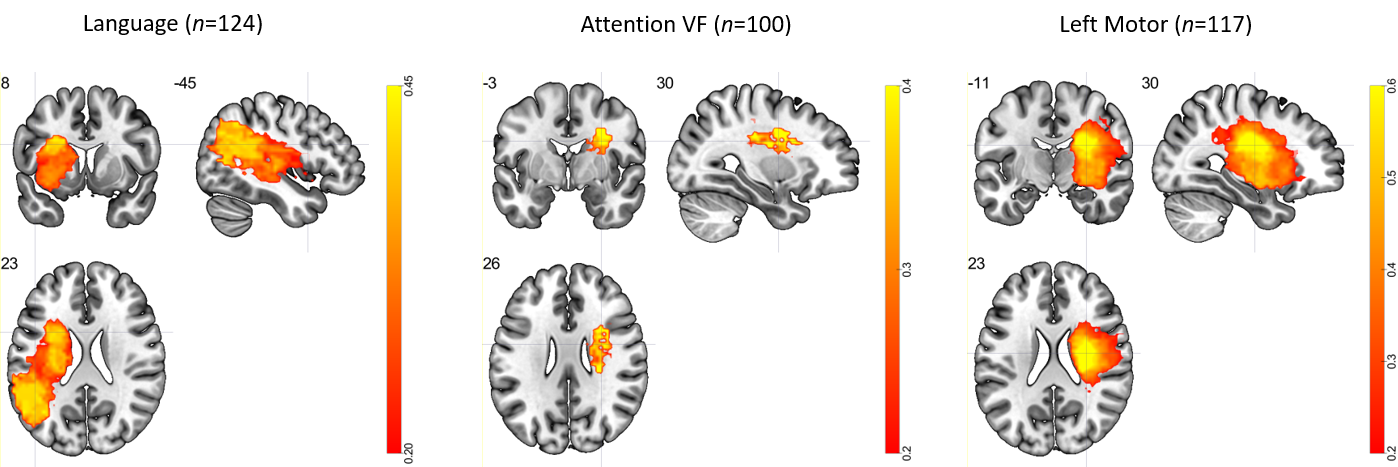

**Supplementary Figure 4.** Results of voxel-based lesion analyses. Mass univariate point-biserial correlation analyses were performed to relate voxel-level lesion statuses to each residual behavioral measure (i.e. after regressing lesion volume out of the behavioral measures). Results that survived a false discovery rate threshold of 0.05 are shown for each analysis.

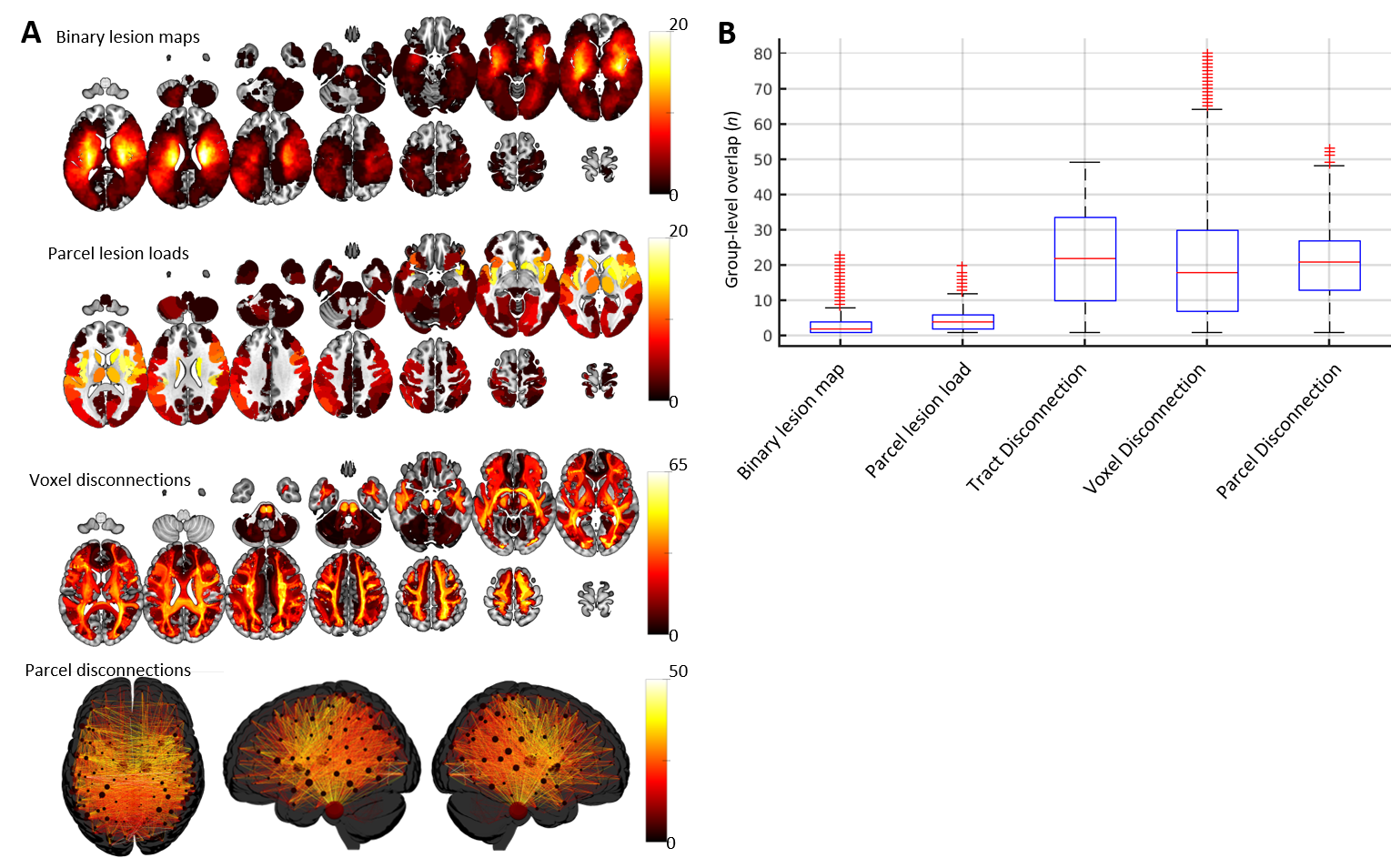

**Supplementary Figure 5.** Damage and disconnection frequencies. **A.** Group-level (*n*=132) damage and disconnection frequencies are shown for binary lesion maps, parcel lesion loads, voxel disconnections, and parcel disconnections. For the latter three measures, only parcels/voxels/connections with at least 10% damage/disconnection were included for each patient. **B.** Values from the maps shown in (A) are plotted as boxplots to illustrate how the measures produced by the toolkit can enable identification of common structures affected among patients with non-overlapping lesions. The boxplots show the distributions of group-level overlaps (i.e. number of patients with damage or disconnection – y-axis) across elements (i.e. voxels, parcels, tracts, etc.) from each measure (x-axis). While parcel-level damage frequencies (i.e. parcel lesion loads) tends to be higher than voxel-level damage frequencies (i.e. binary lesion maps), disconnection frequencies (i.e. remaining three measures) tend to be much higher than either of these. This illustrates how summarizing lesion data in terms of anatomical structures, especially white matter connections, can highlight common structural damage among patients with relatively low voxel-level lesion overlap.

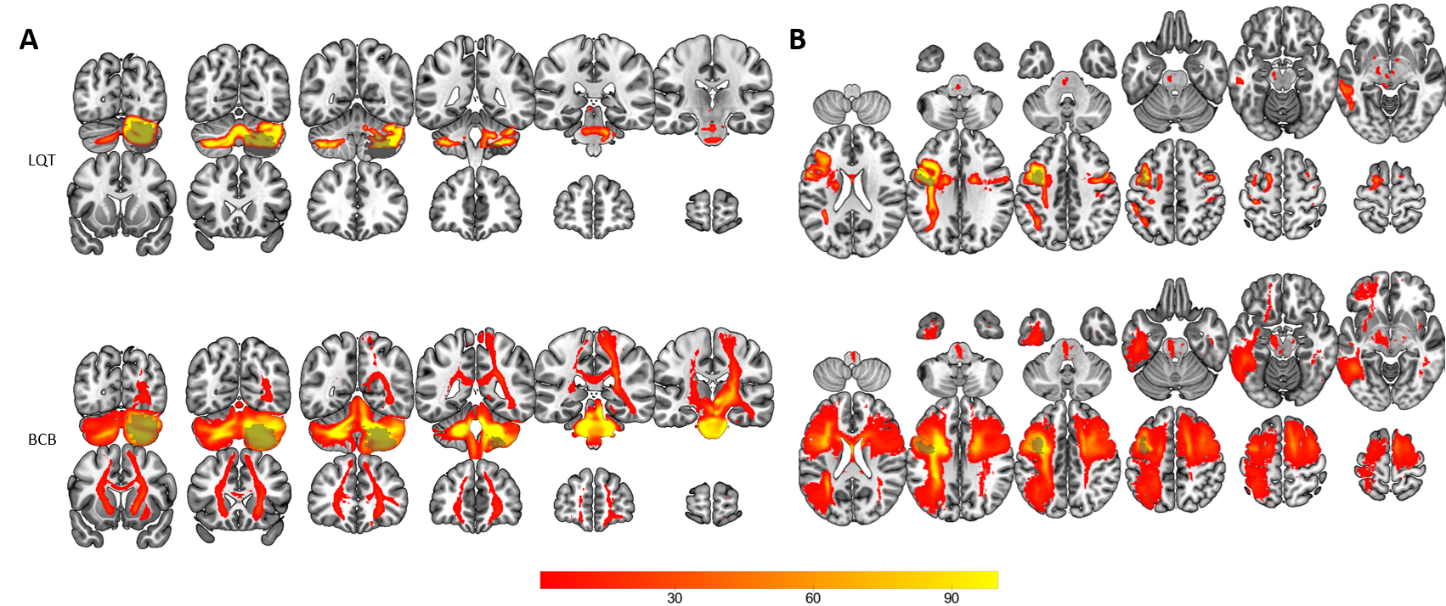

**Supplementary Figure 6.** Qualitative comparison of voxel-wise disconnection maps produced by the Lesion Quantification Toolkit (LQT) with disconnection maps produced by the Brain Connectivity and Behavior (BCB) toolkit. **A**. Disconnection severity map produced by LQT (top) and disconnection “probability” map produced by BCB for a patient with a right cerebellar lesion (transparent grey overlay). Note that while the map produced by LQT (top) features voxels consistent with intra-cerebellar disconnections and disconnections between the cerebellum and brainstem, the map produced by BCB (bottom) additionally features voxels in the cerebral white matter (including corpus callosum) that are suggestive of cortical disconnections. The latter observations are suggestive of fiber tracking errors in the healthy reference group utilized by BCB. **B.** Disconnection severity map produced by LQT (top) and disconnection “probability” map produced by BCB for a patient with a left lateral prefrontal lesion (transparent grey overlay). Note that while the map produced by LQT (top) only features voxels consistent with disconnections of left fronto-parietal/fronto-temporal, homotopic, and left cortico-subcortical projection pathways, the map produced by BCB (bottom) additionally features voxels in the right hemisphere that seem consistent with contralateral fronto-parietal disconnections (i.e. 2^nd^ and 3^rd^ slices from the left, bottom row), again suggestive of fiber tracking errors in the healthy reference group utilized by BCB. While qualitative, these results highlight the potential for methods based on un-vetted fiber tracking in healthy reference groups to produce biologically implausible disconnection results, although detailed quantitative comparisons of different white matter disconnection measures are necessary to allow for strong conclusions about which measures are most useful in practice. BCB maps were created according to the procedure described by Salvalaggio et al., (2020).
