## Supplementary material for "Lesion Quantification Toolkit: A MATLAB software tool for estimating grey matter damage and white matter disconnections in patients with focal brain lesions": User Manual

**User Manual**

© 2020, Joseph C. Griffis, PhD, Washington University in St. Louis Department of Neurology

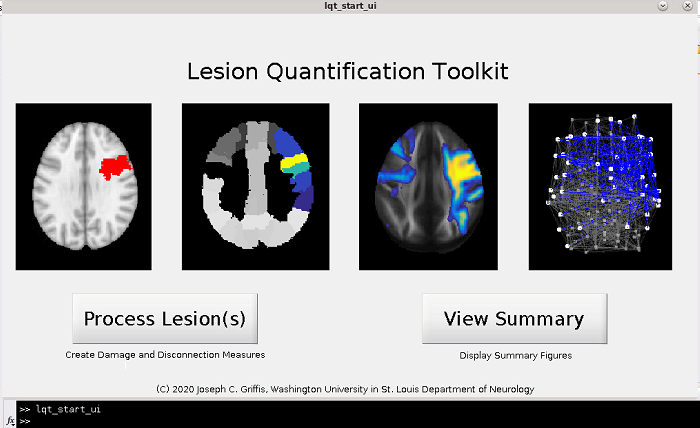

**1. Requirements**

**1a. Software Requirements**

This toolkit runs in MATLAB (it has only been tested in R2015a) and requires the DSI_Studio software package (available at <http://dsi-studio.labsolver.org/>). It has only been tested in a Linux environment.

The following MATLAB toolboxes are required, and are included in the toolkit:

1. matlab_nifti (<https://www.mathworks.com/matlabcentral/fileexchange/8797-tools-for-nifti-and-analyze-image>)

2. Brain Connectivity Toolbox (<https://sites.google.com/site/bctnet/>)

3. GRETNA (<https://www.nitrc.org/projects/gretna/>)

The toolkit also requires the HCP842 population average diffusion template and tractography atlas described in [1], which are included in the toolkit, but are also available at <http://brain.labsolver.org/>.

**1b. Other Requirements (Inputs)**

**Lesion files:** This toolkit does not perform lesion segmentation or anatomical registrations, and requires that the user supply MNI-registered binary lesion segmentation files in the *nifti* file format (i.e. .nii, .nii.gz).

For compatibility with the included parcellation files, it is necessary that the lesion files have 1mm isotropic voxel dimensions and image dimensions of 182x218x182. These are the same dimensions as the MNI_152_T1_1mm.nii.gz file contained in /Lesion_Quantification_Toolkit/Support_Tools.

Regardless of image dimensions, it is necessary that the lesion files be registered to MNI template space prior to using the toolkit in order to avoid errors in processing.

**Parcellation files:** Multi-resolution brain parcellation files are included with the toolkit. They are located in /Lesion_Quantification_Toolkit/Parcellations/Schaefer_Yeo, and correspond to the cortical resting-state functional parcellations described in [2]. (<https://pubmed.ncbi.nlm.nih.gov/28981612/>). Since these parcellations only cover the cortex, modified versions were created by adding subcortical/cerebellar parcels from the AAL atlas [3] and a brainstem parcel from the Harvard-Oxford Subcortical atlas (available at <https://fsl.fmrib.ox.ac.uk/fsl/fslwiki/Atlases>) to allow for complete brain coverage. The combined parcellations are located in /Lesion_Quantification_Toolkit/Parcellations/Schaefer_Yeo/Plus_Subcort. These parcellations are in MNI template space with 1mm isotropic voxel dimensions and image dimensions of 182x218x182. In order to use these files, it is necessary to ensure that the lesion files have identical voxel and image dimensions. User-supplied parcellations may also be used, but they must be registered to the MNI template space and have identical voxel and image dimensions to the lesion files. For each included parcellation file, information about the parcel labels, co-ordinates, and network assignments can be found in a MATLAB table contained within a corresponding .mat file that has the same name as the parcellation. This information can be used to, for example, summarize parcel-level damage and disconnection measures in terms of resting-state network assignments.

**2. How to use the toolkit**

Note: The toolkit can be used via either a graphical user interface (GUI) or by running simple MATLAB scripts. The script-based approach is recommended since it is faster and can improve reproducibility. If you plan to use the toolkit via the graphical user interface frequently (GUI), then prior to using the toolkit, then you should create a ‘dsi_path.mat’ file containing the filepath to the dsi_studio executable file (saved as a string variable named “dsi_path”) in the main Lesion_Quantification_Toolkit folder prior to running the GUI. If you don’t create this file, then you will be prompted to select the dsi_studio executable file the first time that you run the GUI, and this file will be automatically saved for future use.

**2a. Graphical User Interface**

The toolkit features a basic graphical user interface (GUI) that allows the user to create measures for single patients or for groups of patients (i.e. batch processing), and also allows the user to review summary figures for previously processed patients.

**To use the GUI to process a new lesion or set of lesions**:

1. Open MATLAB and navigate to the directory containing the Leson_Quantification_Toolkit folder.

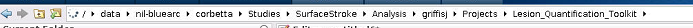

2. Right click on the “Functions” folder, and select “Add to path”.
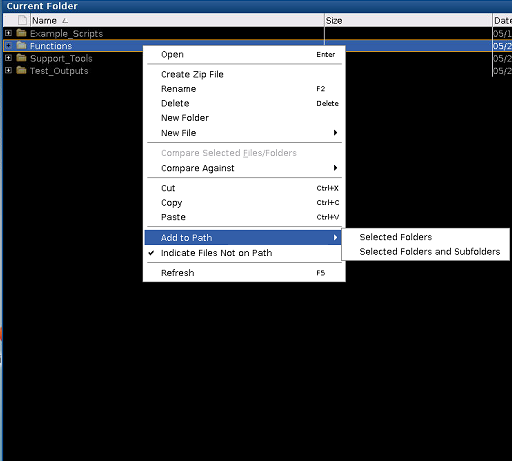

3. It is recommended that you create a “dsi_path.mat” variable containing the path to the DSI_studio executable file in the Lesion_Quantification_Toolkit directory prior to running the GUI.

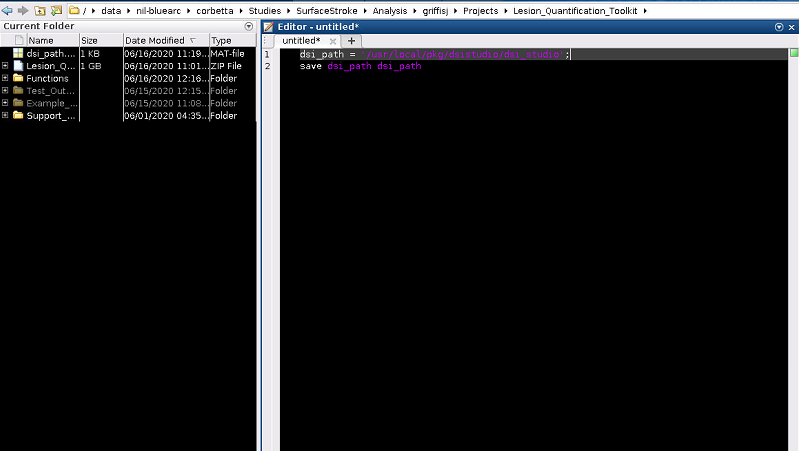

4. If you do not create this file prior to running the GUI, then you will have to manually navigate to and select the DSI_studio executable file the first time that you run the GUI. However, the path to the DSI_studio file will then be automatically saved for future use.

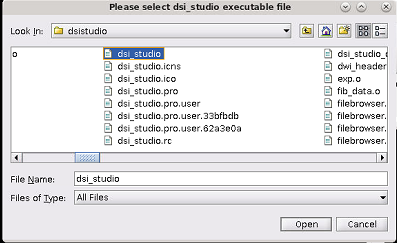

5. In the MATLAB command window, type “lqt_start_ui” (without quotes). This will start the GUI. From the GUI window, select “Process Lesion(s)” to create damage and disconnection measures using lesion files from a single patient or group of patients.

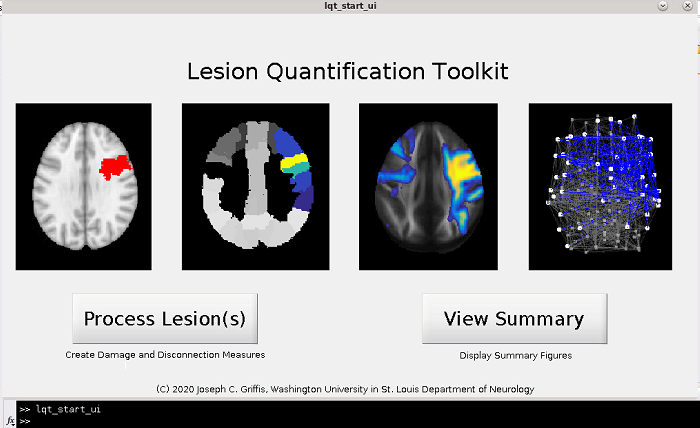

6. Using the interactive file explorer, select the lesion file(s) that you wish to process. These must be *nifti* files (.nii, .nii.gz) that are registered to MNI space and that have identical voxel and image dimensions to the brain parcellation you wish to use. You may select multiple lesion files. If multiple lesion files are selected, then they will be treated as separate patients, and separate subfolders will be created within the output folder (selected during the previous step) that contain the outputs for each lesion. This allows for batch processing of lesions from groups of patients.

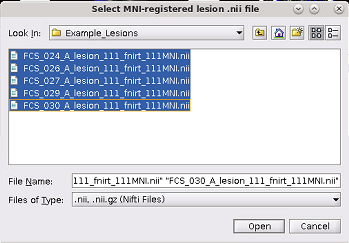

7. After you have selected your lesion file(s), you will need to select your parcellation file. A set of multi-resolution resting-state parcellations by Schaefer et al., (2018), which have been expanded to include additional subcortical and cerebellar parcels from the AAL and Harvard-Oxford anatomical atlases are included in the toolkit as default parcellations. A question box will ask you if you want you use a default parcellation. If you select “Yes”, then a file selection box will open, and you will be able to select the desired parcellation from the list. Note that to use a default parcellation, your lesion file must have dimensions of 182x218x182. If you select “Yes”, then your output files will be automatically suffixed with information about the selected parcellation (e.g. Schaefer_1000_17). If you select “No”, then a file explorer window will open, and you can use this window to select the desired parcellation. Your selected parcellation file must be a *nifti* file (i.e. .nii or .nii.gz) that is registered to MNI space and that has identical voxel and image dimensions to the lesion files. You may only select a single brain parcellation file. If you choose to select a parcellation other than the default parcellation, you will be asked to provide a file suffix that will be appended to the output files.

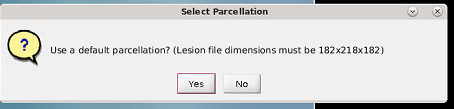

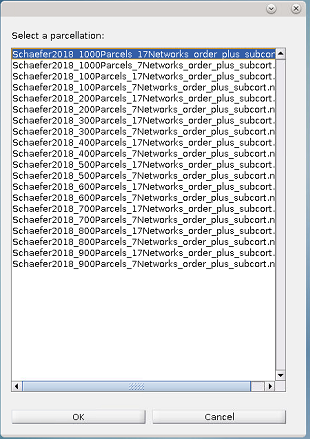

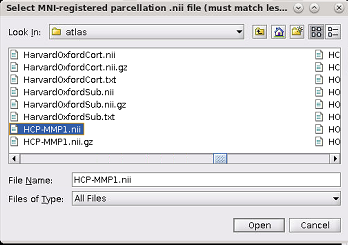

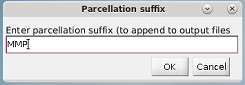

8. Next, you will be prompted to select the folder where you wish to save your output files. Using the interactive file explorer, select the folder where you wish to store the outputs. Within this folder, an “Atlas” folder will be created that will include reference files defined based on the HCP-842 tractography atlas and your chosen parcellation, and new subfolders will be also be created that will contain the outputs for each patient.

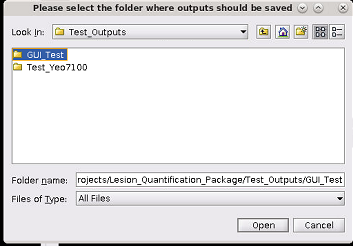

9. Next, a dialog box will appear. You will also be asked to specify the connectivity type. The default is “end”, which defines the connections between two parcels as those streamlines that end in both parcels. The alternative option is “pass”, which defines the connections between two parcels as those streamlines that either end in or pass through both parcels. End is recommended but will produce sparser connectivity matrices.

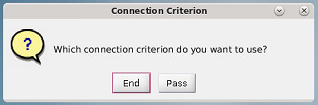

10. Finally, you will also be asked to input the % spared binarization threshold for connections to include in the calculation of structural shortest path lengths (SSPLs). This option defines the minimum % of streamlines connecting two parcels that must be spared for the connection between those parcels to be considered as spared in the calculation of parcel-wise SSPLs. The default is 50, meaning that parcel pairs that suffered greater than 50% disconnection are not included in the calculation of SSPLs. You will also be asked to input a smoothing kernel (any real number) which will be used to apply Gaussian smoothing to the voxel percent disconnection maps.

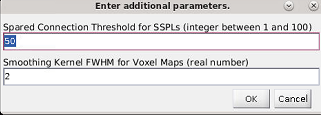

Once the required information has been provided, the damage and disconnection measures will be automatically created and output into the subject subfolders located within the output directory. The outputs are described in subsequent sections.

**How to view summary figures after processing:**

To create and view summary figures after processing, click on the “View Summary Figures” button on the GUI menu. Then, navigate to the subfolder of the patient for whom you wish to create summary figures (located in the output directory), and select the cfg.mat file using the interactive file explorer.

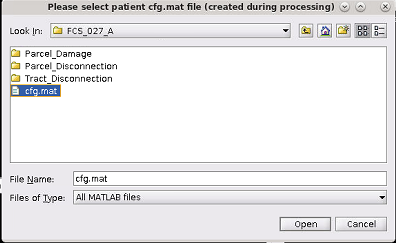

This will create several summary figures:

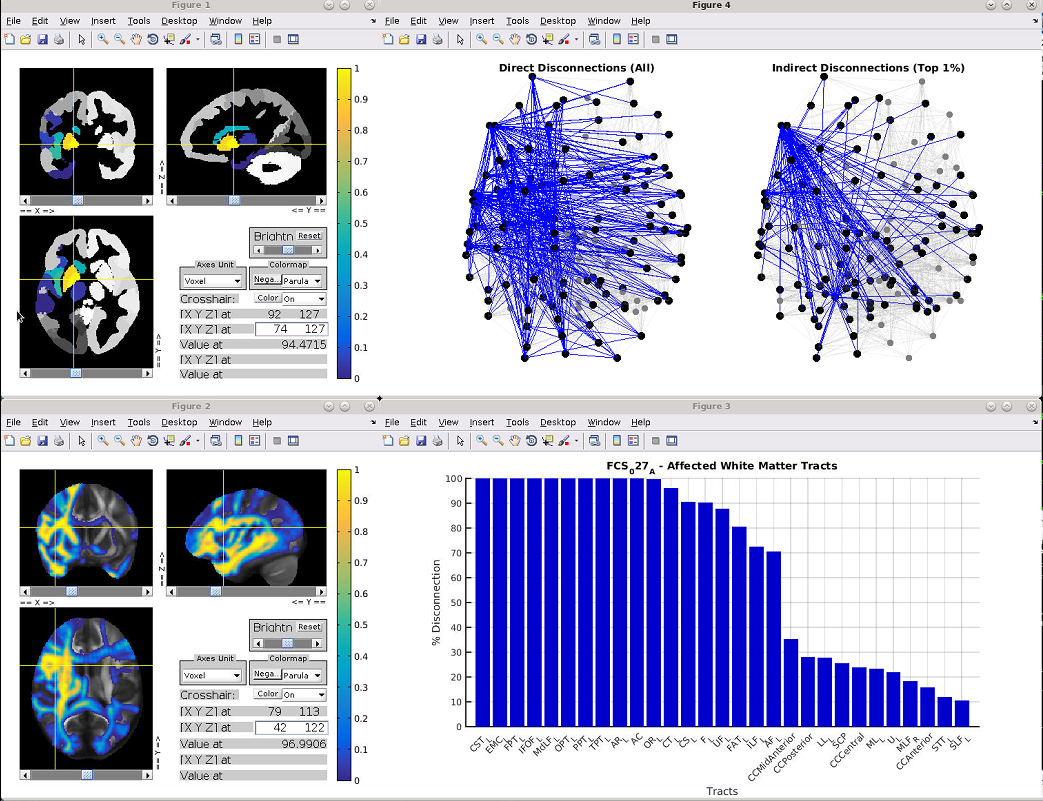

The top left panel (Figure 1) is an interactive viewer that displays the parcel damage map (see Section 3a). The color scale indicates the proportion of voxels within each parcel that were damaged by the lesion. The bottom left panel (Figure 2) is an interactive viewer that displays a TDI percent disconnected fiber map (see Section 3c). The color scale indicates the proportion of streamlines within each voxel that were disconnected by the lesion. The bottom right panel (Figure 3) shows the percent of streamlines associated with each fiber tract that were disconnected by the lesion, sorted in descending order. The top right panel (Figure 4) shows a graph brain representation of the parcel-wise direct disconnections associated with the lesion (left plot – Direct Disconnections (All)) and an analogous representation of the top 1% of indirect disconnections associated with the lesion (see Section 3c). Graph brain representations are by default viewed from top-down.

**2b. MATLAB Scripts**

MATLAB scripts provide a more efficient and flexible way of using this toolkit. Example scripts with detailed comments are included for both single-patient and multi-patient (i.e. batch) processing in /Lesion_Quantification_Toolkit/Example_Scripts. These scripts are intended to be easily modifiable, and they provide a template for producing all of the outputs.

**To set up a script:**

First, add /Lesion_Quantification_Toolkit/Functions and /Lesion_Quantification_Toolkit/Support_Tools (all subfolders) to the MATLAB path.

Next, set up the configuration (cfg) structure. This structure will contain the file paths, user options, and parameter choices. The user should set the following fields:

1. **cfg.dsi_path** – this should contain the full path to the dsi_studio program.
2. **cfg.source_path** – this should contain the full path to the folder containing the HCP842.trk.gz file and tractography atlas files (i.e. …/Lesion_Quantification_Package/Support_Tools/Tractography_Atlas/)
3. **cfg.out_path** – this should contain the full path to the folder where results will be output. Note that subject specific sub-folders will be created within this folder.
4. **cfg.lesion_path** – this should contain the full path to the MNI-registered lesion *nifti* file. For batches, this can be updated in a loop that iterates over patients.
5. **cfg.parcel_path** – this should contain the full path to the MNI-registered brain parcellation *nifti* file. This file should have identical voxel and image dimensions as the lesion file.
6. **cfg.pat_id** – this should contain a patient identifier (e.g. “patient_01”), which will be used to create the subject-specific subfolders within the directory supplied to cfg.out_path as well as to prefix the output files for each patient. For batches, this can also be updated in a loop that iterates over patients.
7. **cfg.file_suffix** – this should contain an identifier for the processing run (e.g. the parcellation used), which will be appended to the output files within each subject subfolder.
8. **cfg.con_type** – this will specify the criteria for defining structural connections. There are two options: “end”, which defines the connections between two parcels as those streamlines that end in both parcels, or “pass”, which defines the connections between two parcels as those streamlines that either end in or pass through both parcels. “End” is recommended but will produce sparser connectivity matrices.
9. **cfg.sspl_spared_thresh** – this will specify the minimum percent of streamlines connecting two parcels that must be spared for the connection to be considered when computing SSPLs from the patient spared connectivity matrix. The default is 100, meaning that only parcel pairs whose structural connections were completely spared by the lesion will be considered in the SSPL computation (see section 3c).
10. **cfg.node_label** – this is an optional field that will specify the names for each node (i.e. region, parcel) in the brain parcellation. It should be an n_regions-by-1 cell array of strings corresponding to node names.
11. **cfg.node_color** – this is an optional field that will specify different color levels for each node in the brain parcellation when viewed in external viewers. It should be an n_region-by-1 array of integer values, where each unique value corresponds to a distinct region identifier (e.g. resting-state network assignment).
12. **cfg.show_summary_figs** – this field may be set to either 0 or 1. If set to 0, no summary figures will be displayed. If set to 1, then a summary figure will be created after processing is complete.
13. **cfg.parcel_coords** – this is an optional field can be set to an n_regions-by-[x,y,z] matrix containing the MNI coordinates for each parcel in the brain parcellation, which will be used when creating files for external viewing and for creating the summary ball-and-stick brain graph visualization if cfg.show_summary_figs==1. If it is empty, then parcel coordinates will be estimated from the brain parcellation file.
14. **cfg.delta_sspl_thresh** – this field can be set to a percentile threshold for displaying SSPL increases in the summary figures if cfg.show_summary_figs==1. For example, if it is set to 99, then only the top 1% of SSPL increases will be displayed in the summary figure.
15. **cfg.parcel_dmg_thresh** – this field can be set to a percent damage threshold for removing regions from the SSPL increase summary figure. For example, if it is set to 10, then it will not show SSPL increases involving parcels with >10% damage.
16. **cfg.tract_sdc_thresh** – this field can be set to a percent disconnection threshold for showing tract disconnections in the summary figure if cfg.show_summary_figs==1. For example, if it is set to 5, then the tract disconnection summary figure will only show data for tracts with >5% disconnection.

Next, call the following functions using the cfg variable as an input argument:

1. util_get_parcel_damage(cfg)
2. util_get_tract_discon(cfg)
3. util_get_parcel_cons(cfg)

If cfg.show_summary==1, then to view summary figures after processing is complete, also call: util_plot_summaries(cfg). This will produce summary figures of the damage and disconnection measures for this patient.

**3. Outputs**

This toolkit is intended to facilitate the quantification of the local (e.g. regional damage) and distributed (e.g. inter-regional disconnections) structural consequences of focal brain lesions (e.g. strokes, tumors, etc.) for research purposes. Disconnection measures are based on the curated population-level structural connectome described in [1]. The damage and disconnection measures created by this toolkit largely correspond to the measures described in [4], [5], which are also included in /Lesion_Quantification_Toolkit/Publications.

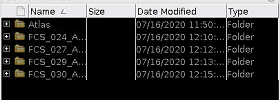

**3a. Atlas** **Folder**

Within the output directory, a folder titled “Atlas” will be created. This folder will contain several measures created using the HCP-842 tractography atlas in conjunction with the selected brain parcellation. The measures contained within this folder are used in the computation of the patient disconnection measures:

1. **Aggregate streamline map** – a .trk file containing the aggregate streamlines from the HCP-842 atlas.
2. **Voxel-wise tract density map** – a .nii file containing the tract density image (TDI) of the aggregate streamline map.
3. **Atlas structural connectivity matrix** – a .mat file containing the variable “connectivity”, which is an adjacency matrix containing the atlas structural connectivity matrix for the user-selected brain parcellation.

**3b. Patient Folders**

Within the output directory, subfolders will be created for each patient. Within each patient subfolder, the damage and disconnection measures will be output to the following subfolders:

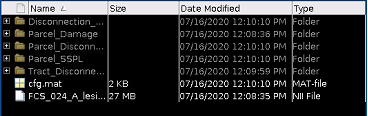

**Parcel_Damage**

For a given lesion and brain parcellation, the parcel damage measures quantify the amount of damage sustained by each brain parcel (i.e. region). This measure is computed by overlapping the lesion with each parcel and computing the percent of voxels within the parcel that overlap with the lesion. Depending on the type of brain parcellation used, this provides a straightforward way of reducing the dimensionality of the lesion by summarizing it in terms of the damage sustained by anatomically- and/or functionally-defined brain regions, rather than in terms of individual voxels.

1. **Parcel damage map** – a .nii file where the voxels within each parcel from the supplied parcellation are assigned values corresponding to the percent of voxels from that parcel that were affected by the lesion.
2. **Parcel damage array** – a .mat file containing a 1 x n_parcels array, where the value for each parcel index corresponds to the percent of voxels from that parcel that were affected by the lesion.

**Tract_Disconnection**

For a given lesion, the tract disconnection measures quantify the severity of disconnection sustained by each of 70 canonical white matter tracts. These tracts correspond to the 66 tract segmentations reported in [3] (see /Lesion_Quantification_Toolkit/Publications), with the exception that the Corpus Callosum was split into 5 segments based on the FreesurferSeg corpus callosum segment ROIs, resulting in 70 tracts.

The tract disconnection measure is computed by embedding the lesion into the HCP842 tractography atlas, and then measuring the percent of streamlines within each tract that intersect the lesion. This provides a straightforward way to estimate the severity of tract-level white matter disconnections caused by a lesion, and importantly, provides a measure of *disconnection* *severity* rather than *tract overlap* (i.e. a small lesion that intersects all streamlines contained within a tract will result in 100% disconnection, even though it may only overlap with a small section of the tract).

1. **Tract disconnection array** – a .mat file containing a 1x70 array, where the value for each tract index corresponds to the percent of streamlines from that tract that intersected the lesion. Tract names are also provided in a 1x70 cell array for reference.

**Parcel_Disconnection**

**Direct parcel disconnection files:** For a given lesion and brain parcellation, the parcel disconnection measures quantify the severity of the expected lesion-induced structural disconnections between pairs of parcels (i.e. brain regions) with direct structural connections in the atlas-based structural connectome. These measures are computed by first constructing a parcel-wise structural connectome using the user-selected brain parcellation and the HCP842 tractography atlas (this will be output in a subfolder within the output director named “Atlas”), embedding the lesion into the atlas-derived parcel-wise structural connectome, and identifying the parcel-wise connections that intersect the lesion. These measures provide insights into the direct effects of the lesion on the structural connectome.

1. **Raw parcel disconnection matrix** - a .mat file with the suffix .connectivity.mat. This contains both the raw structural disconnection matrix (connectivity) and parcel names (name), although you must use char(name) to convert to actual names. Values in this disconnection matrix correspond to the number of streamlines connecting each region pair that intersected the lesion. Because raw streamline counts vary substantially across region pairs, this matrix is not very interpretable. This matrix is primarily used to create the percent parcel disconnection matrix and the percent spared connection matrix.
2. **Percent parcel disconnection matrix** - a .mat file with the suffix _percent_parcel_SDC.mat. This file contains a disconnection adjacency matrix where each cell quantifies the percent of streamlines connecting a region pair in the atlas-based structural connectome that intersected the lesion, and therefore provides an interpretable measure of disconnection severity for each edge (i.e. connection, region pair) in the structural connectome. This matrix can be used in connectome-based lesion-symptom mapping (CLSM)-style analyses [4], [6].
3. **Percent spared parcel connectivity matrix** - a .mat file with the suffix _percent_parcel_spared_SC.mat. This file contains a spared connection adjacency matrix where each cell quantifies the percent of streamlines connecting each region pair in the atlas-based structural connectome that were spared by the lesion (i.e. it is equal to 100 – the parcel direct disconnection matrix). This is used to identify increases in structural shortest path lengths (SSPLs) associated with the lesion.
4. **Disconnection connectogram (for external viewing)** - a .txt file with the suffix .connectogram.txt. This file contains a connectomgram that can be viewed on http://mkweb.bcgsc.ca/tableviewer/visualize/ by checking the two size options in step 2A (col with row size, row with col size)
5. **Disconnection .node file (for external viewing)** - a .node file. This file contains the node information for external connectome viewers (e.g. MRIcroGL). Node sizes are proportional to the severity of disconnections for each node. Node colors can be pre-assigned in the .cfg file (cfg.node_color), but if not, they will be analogous to node size.
6. **Disconnection .edge file (for external viewing)** – a .edge file with the suffix _percent_parcel_SDC.edge. This file contains the percent SDC matrix in a format that can be loaded into external viewers (e.g. MRICroGL).
7. **Spared connection .edge file (for external viewing)** - a .edge file with the suffix _percent_parcel_spared_SC.edge. This file contains the percent spared SC matrix in a format that can be loaded into external viewers (e.g. MRICroGL).
8. **Spared connection .node file (for external viewing)** – a .node file with the suffix _percent_parcel_spared_SC.node. This file contains the node information for external connectome viewers. Node sizes are proportional to the total spared connections for each node.

**Disconnection_Maps**

**Map files:** The measures output into this folder include several disconnection map files. While they are output into this folder for convenience since they are computed along with the parcel disconnection files, the map files are not dependent on the parcellation as they are simply maps of the streamlines that intersect the embedded lesion.

1. **Streamline disconnected fiber map** - a .trk.gz file. This contains all of the streamlines that intersected the lesion and can be viewed using e.g. DSI_Studio. This allows for high-detail visualizations of the disconnections associated with a lesion as streamlines.
2. **Tract density imaging (TDI) disconnected fiber map** -. a .tdi.nii.gz file named the same way as the .trk.gz file. This contains a nifti image volume with track density imaging (TDI) values from the .trk.gz file at each voxel. It is essentially a way of converting the .trk.gz file into voxel space. Higher values indicate higher streamline densities at each grid element (voxel). This map is primarily used for creating the percent disconnected fiber map.
3. **TDI percent disconnected fiber map** - a .nii file with the suffix _percent_tdi.nii. For each voxel, values correspond the % reduction in streamline density relative to the atlas when accounting for the effects of the lesion. These files are derived from the TDI disconnected fiber map, and allow for high-detail visualizations of the disconnections associated with a lesion as a voxel-wise map. These maps can also be used in lesion-symptom mapping or voxel-based morphometry-style analyses.

**Parcel_SSPL**

**Higher-order disconnection files:** For a given lesion and brain parcellation, the higher-order parcel disconnection measures quantify expected lesion-induced increases in the topological distances, defined in terms of structural shortest path lengths (SSPLs), between pairs of parcels (i.e. brain regions) with and without direct structural connections in the atlas-based structural connectome [7]. SSPLs correspond to the minimum number of direct structural connections that must be traversed to establish a structural path between a given pair of parcels (i.e. brain regions). These measures are computed by first computing an atlas-based SSPL matrix by applying a breadth-first search algorithm to the binarized atlas-based structural connectome, computing a patient-specific SSPL matrix by applying the same algorithm to the binarized patient spared SC matrix, and identifying pairs of parcels (i.e. brain regions) with longer SSPLs in the patient-specific SSPL matrix than in the atlas-based SSPL matrix [5]. This measure, as computed here, depends on the binarization threshold applied to the percent spared parcel connectivity matrix (set in **cfg.sspl_spared_thresh**). These measures provide insights into how the structural network topology is likely to be altered by the lesion.

1. **Raw parcel structural shortest path length (SSPL) matrix** - a file with the suffix _SSPL_matrix.mat -- this contains the raw SSPL matrix computed from the binarized spared SC matrix using the breadth-first search algorithm. Values within each cell correspond to the minimum number of structural connections that must be traversed to establish a path between each parcel pair. For example, parcel pairs with direct structural connections will have SSPLs equal to 1, while parcel pairs that lack direct structural connections with each other but that both have direct structural connections with a third parcel will have SSPLs equal to 2.
2. **Delta parcel SSPL matrix** - a file with the suffix _delta_SSPL_matrix.mat -- this contains the expected lesion-induced increases in SSPLs relative to the atlas SSPL matrix. In previous work, we have considered increases in SSPLs between regions that lack direct structural connections to be “indirect disconnections” [5]. Note: both direct and “indirect” disconnections are contained in this matrix, but direct disconnections can be removed by masking out cells contained within the direct parcel disconnection matrix.
3. **Delta SSPL .node file (for external viewing)** – a .node file with the suffix _delta_SSPL.node -- this contains the node information for loading the delta SSPL matrix into an external viewer. Node sizes are proportional to the SSPL increases for each node. Node colors can be set using cfg.node_color, but if not, are analogous to node sizes.
4. **Delta SSPL .edge file (for external viewing)** – a .edge file with the suffix _delta_SSPL.edge. This file contains the percent SDC matrix in a format that can be loaded into external viewers (e.g. MRICroGL).
5. **Indirect SDC matrix** – a .mat file with the suffix _indirect_SDC_matrix.mat. This file contains the delta SSPL matrix with cells corresponding to direct disconnections (i.e. non-zero cells in the parcel direct disconnection matrix) masked out, and therefore only including increases for parcel pairs that did not sustain a direct disconnection.
6. **Indirect SDC .node file (for external viewing)** – analogous to the above .node files.
7. **Indirect SDC .edge file (for external viewing)** – analogous to the above .edge files.

Note: in general, node summary information can be extracted from the .node files by reading them into MATLAB and extracting the appropriate columns.

Note: A copy of the lesion file is also saved into the patient-specific subfolder for future reference.

Note: The various disconnection and spared connection matrices described in this section can also be used to create additional measures. For example, the user could use functions included in the Brain Connectivity Toolbox (located in /Lesion_Quantification_Toolkit/Support_Tools) to estimate lesion-induced reductions global efficiency by estimating the global efficiency of the atlas structural connectome and the patient’s spared structural connectome, and subtracting the global efficiency of the spared connectome from that of the atlas connectome. Because of the large number of possible additional measures that could be derived from the measures described in this section, we have left it up to the user to determine if/how they want to compute such additional measures.

**4. External Viewers**

Several output files are generated to allow for the creation of publication-quality visualizations using external image viewers.

**4a. Disconnected fiber streamline map (.trk file)**

This file allows for high-resolution visualizations of the direct structural disconnections associated with a given lesion, and can be viewed using DSI_studio or other software that is compatible with the .trk file format.

To visualize this file in DSI_studio:

1. Open DSI_studio and select “Step T3: Fiber Tracking”

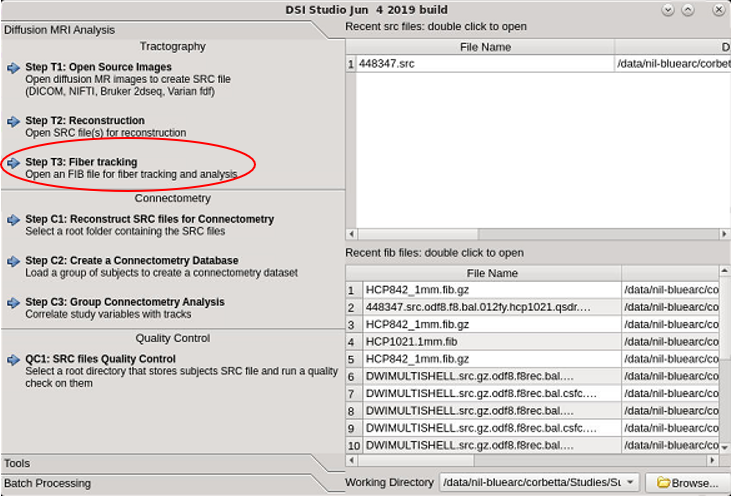

2. Navigate to /Lesion_Quantification_Toolit/Support_Tools/Tractography_Atlas, and select the file “HCP842_1mm.fib.gz”

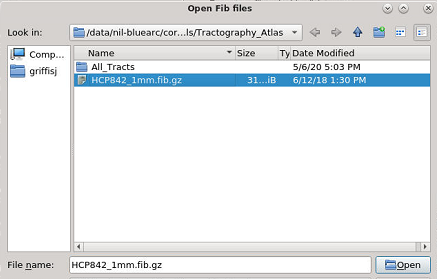

3. From the DSI_Studio interface, select “Tracts” -> “Open Tracts”

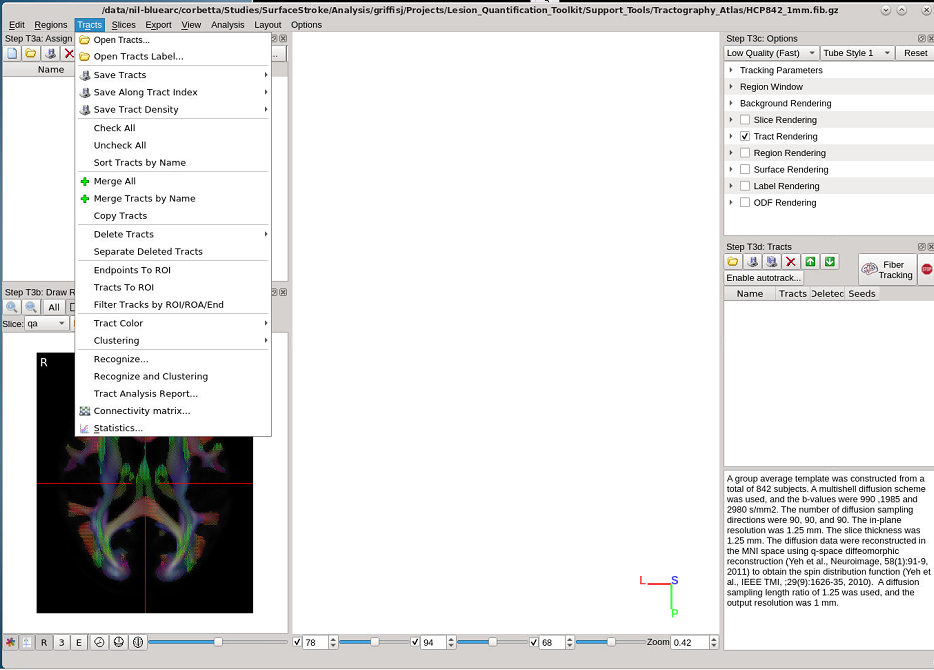

4. Navigate to the “Parcel_Disconnection” subfolder for the patient that you want to visualize, and select the file ending in “.trk.gz”.

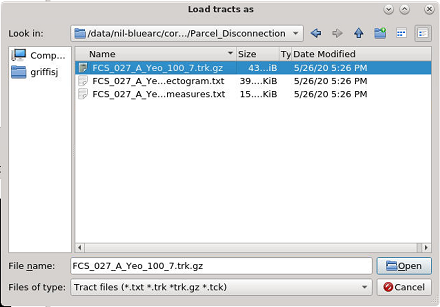

5. Select “Slice Rendering” and the desired underlay image to show brain slices.

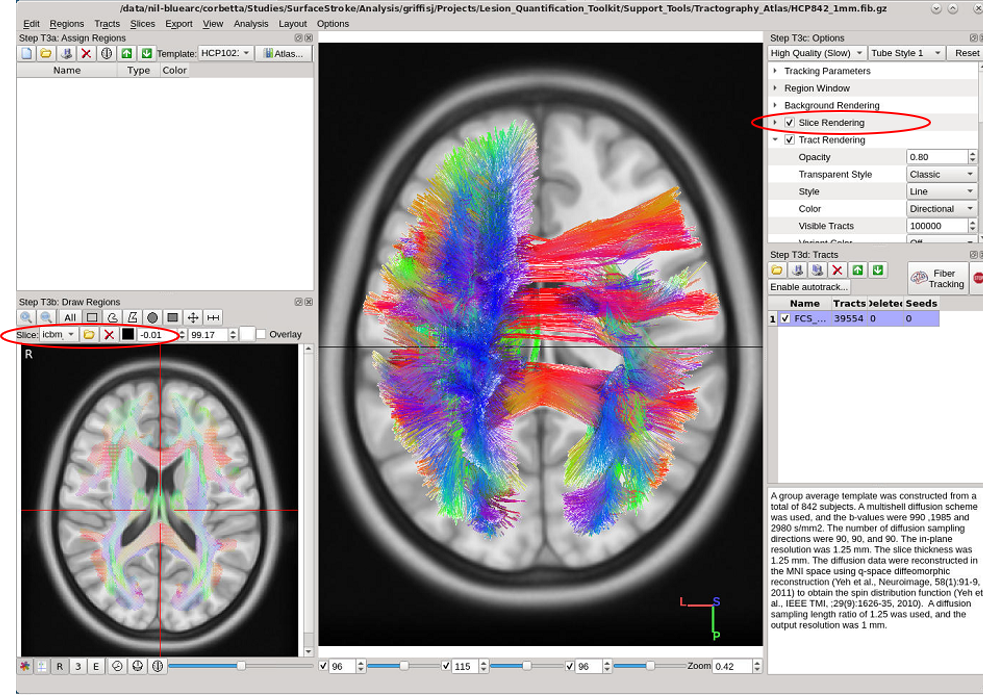

For more details on data visualization in DSI_studio, please consult the DSI_studio manual (<http://dsi-studio.labsolver.org/Manual/tracts-visualization>).

**4b. Visualizing Chord Plot Connectogram**

You can also create chord plot connectogram visualizations of the direct structural disconnection matrix.

To create a chord plot connectogram, first open the following website in a browser:  <http://mkweb.bcgsc.ca/tableviewer/visualize/>.

Then, follow the instructions provided here (under section Step 7: Plot a connectogram): <http://dsi-studio.labsolver.org/Manual/tract-specific-analysis#TOC-STEP-7:-Plot-connectogram>

You will upload the file ending in .connectogram.txt located within the Parcel_Disconnection subfolder.

An example connectogram is shown below:

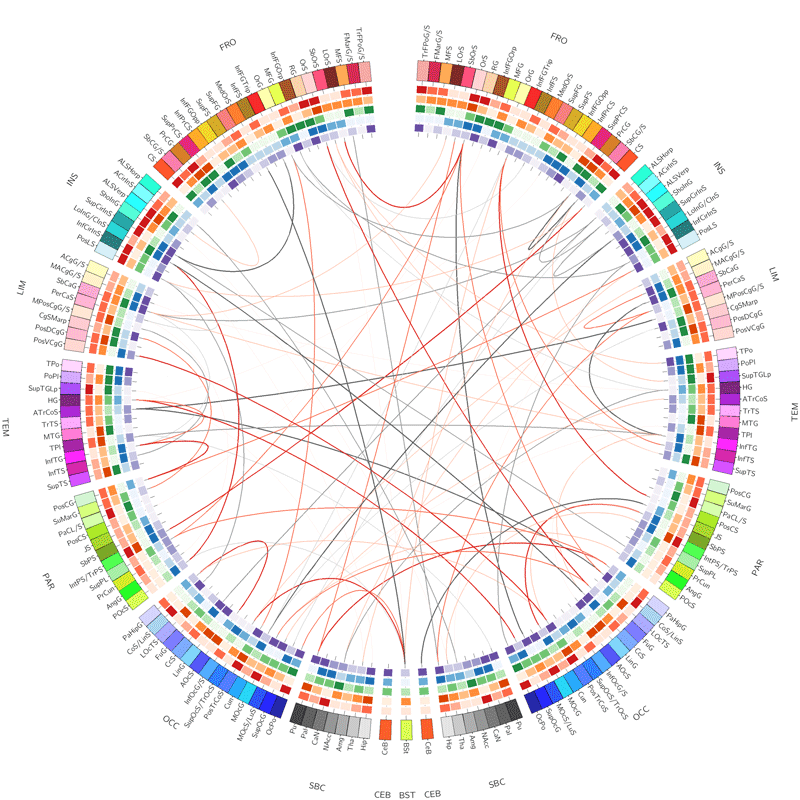

**4c. Visualizing .edge and .node files**

Disconnection matrices are also output as .edge and .node files to allow for ball-and-stick brain network visualizations using external viewer software such as SurfIce (<https://www.nitrc.org/projects/surfice/>).

**To view these files in SurfIce:**

1. Open the SurfIce program and select “Nodes” -> “Add Nodes or Edges” from the menu bar.

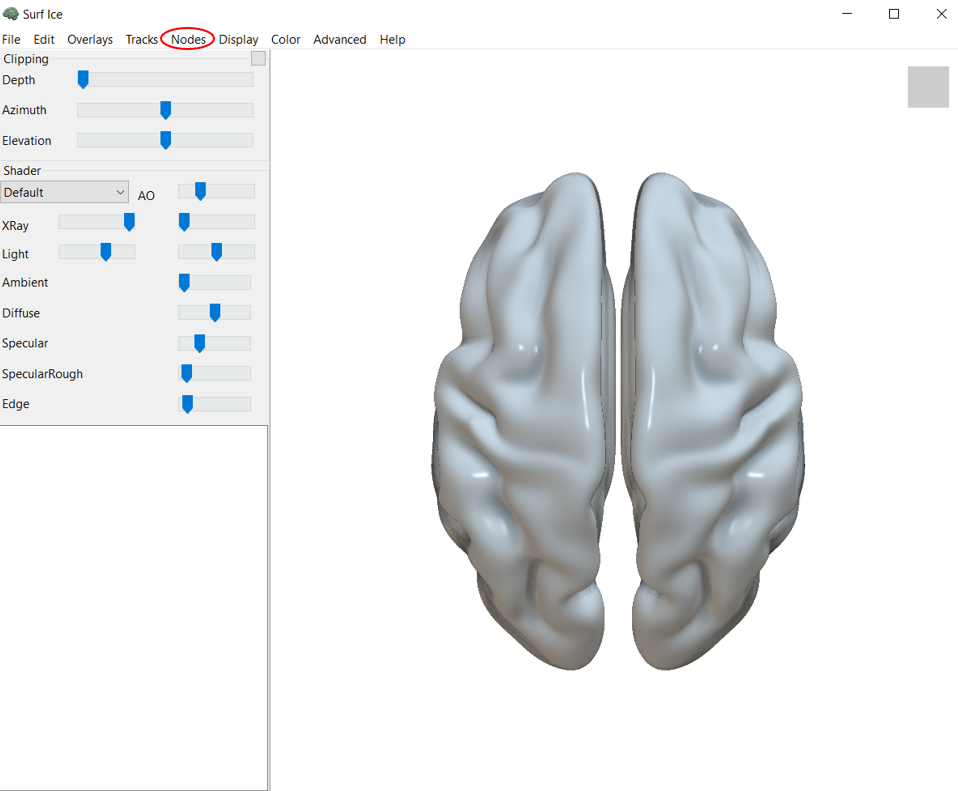

2. Navigate to the “Parcel_Disconnection” subfolder of the patient that you want to visualize, and select the file ending in .node for the desired output. This should also load the corresponding .edge file, but you can also load different .edge files by simply repeating this step and selecting the desired .edge file.

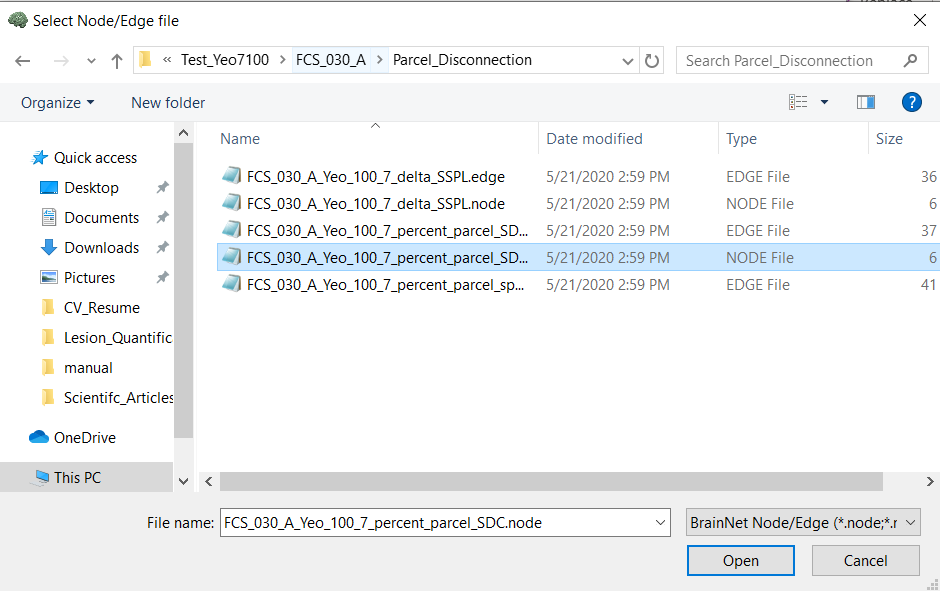

3. Adjust the visualization settings as desired (see <https://www.nitrc.org/projects/surfice/> for more information on visualization settings).

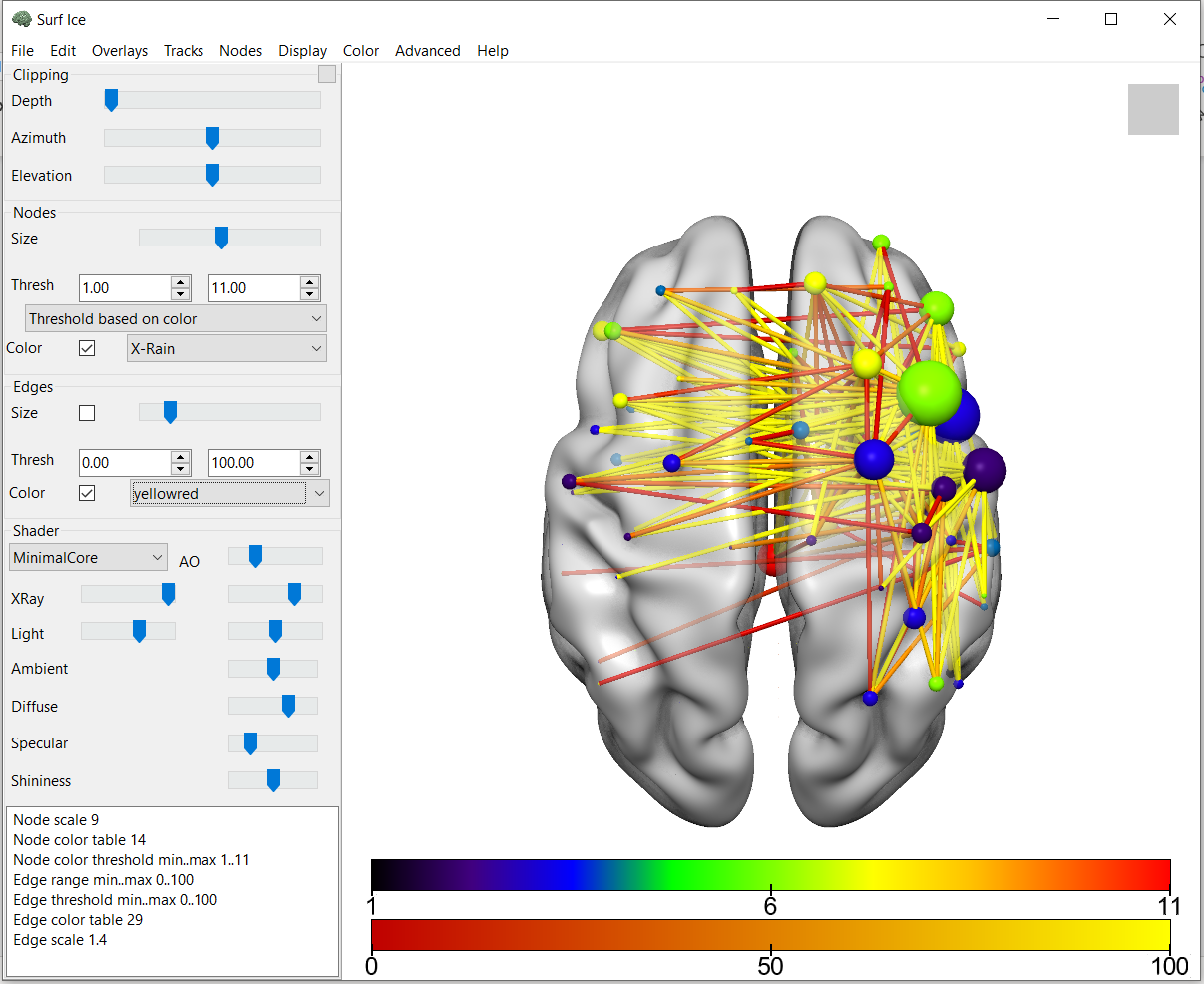

**4c. Visualizing other files**

Other files with the extensions .nii or .nii.gz may also be loaded into any compatible viewing software (e.g. SurfIce, MRIcron, MRIcroGL, FSLview, etc.) for further visualization.
